# “The Heidelberg Five” Personality Dimensions: Genome-wide Associations, Polygenic Risk for Neuroticism, and Psychopathology 20 Years after Assessment

**DOI:** 10.1101/2020.03.23.003889

**Authors:** Urs Heilbronner, Sergi Papiol, Monika Budde, Till F. M. Andlauer, Jana Strohmaier, Fabian Streit, Josef Frank, Franziska Degenhardt, Stefanie Heilmann-Heimbach, Stephanie Witt, Andreas J. Forstner, Adrian Loerbroks, Manfred Amelang, Til Stürmer, Bertram Müller-Myhsok, Markus M. Nöthen, Marcella Rietschel, Thomas G. Schulze

**Affiliations:** Institute of Psychiatric Phenomics and Genomics (IPPG), University Hospital, LMU Munich, Germany; Department of Psychiatry and Psychotherapy, University Hospital, LMU Munich, Germany; Department of Translational Psychiatry, Max Planck Institute of Psychiatry, Munich, Germany; Department of Neurology, Klinikum rechts der Isar, School of Medicine, Technical University of Munich, Germany; Department of Genetic Epidemiology in Psychiatry, Central Institute of Mental Health, Medical Faculty Mannheim, University of Heidelberg, Germany; Institute of Human Genetics, University of Bonn, School of Medicine & University Hospital Bonn, Germany; Department of Child and Adolescent Psychiatry, University Hospital Essen, University of Duisburg-Essen, Essen, Germany; Centre for Human Genetics, University of Marburg, Germany; Department of Biomedicine, University of Basel, Switzerland; Institute of Occupational, Social and Environmental Medicine, Center for Health and Society, Faculty of Medicine, University of Düsseldorf, Germany; Department of Psychology, University of Heidelberg, Germany; Department of Epidemiology, UNC Gillings School of Global Public Health, USA; Department of Psychiatry and Behavioral Sciences, Upstate University Hospital, Syracuse, USA

**Keywords:** Psychoticism, executive function, behavioral control, Type A Behavior, longitudinal cohort study, Locus of Control

## Abstract

The HeiDE study (‘‘Heidelberger Langzeitstudie zu Risikofaktoren und Diagnose chronischer Erkrankungen’’) is a longitudinal population-based study that started in the 1990s and, at baseline, assessed an array of health-related personality questionnaires in 5 133 individuals. Five latent personality dimensions (The Heidelberg Five) were identified and interpreted as Emotional Lability (ELAB), Lack of Behavioral Control (LBCN), Type A Behavior (TYAB), Locus of Control over Disease (LOCC), and Psychoticism (PSYC). A subset of participants (n=3 268; after quality control) were genotyped on whole-genome arrays at follow-up. To further characterize The Heidelberg Five, we analyzed genomic underpinnings, their relations to the genetic basis of the Big Five trait Neuroticism, and longitudinal associations with lifetime psychiatric symptoms. SNP-based heritability was significant for ELAB (34%) and LBCN (29%). Five separate genome-wide association studies (GWAS) using factor scores on personality dimensions as phenotypes were conducted, only the phenotype PSYC yielded a genome-wide significant finding (*p*<5×10^−8^, top SNP rs138223660). Gene-based analyses identified significant findings for ELAB (*Integrin Subunit Beta 5*), TYAB (*Coiled-coil Domain Containing 83*), and PSYC (*Nuclear Receptor Subfamily 1 Group H Member 4*). Polygenic risk scores for Neuroticism, phenotypically related to ELAB, were associated with ELAB, but not with the remaining Heidelberg Five. Longitudinally, all personality dimensions were related to depressive symptoms at follow-up, with ELAB, LBCN, and PSYC also associated with lifetime anxiety symptoms. These results highlight the clinical importance of health-related personality traits, and identify LBCN as a heritable “executive function” personality trait.

## INTRODUCTION

In its widest sense, personality can be conceptualized as “relatively enduring patterns of thoughts, feelings, and behaviors” [1], that constitute hallmarks of individuality. To comprehensively capture all possible manifestations of personality, researchers used the so-called “lexical approach” and analyzed personality-associated words using factor analytic methods, to eventually define the “Big Five” personality traits Extraversion, Neuroticism, Conscientiousness, Agreeableness, and Openness to Experience [2]. These broad dimensions can be used to exhaustively characterize personality and have become the prevailing scientific model. A related but different approach has been to characterize health-related personality dimensions (see [3], [4]), hypothesized to be related to somatic disease such as cardiovascular disease (CVD) and cancer. The HeiDE study (‘‘Heidelberger Langzeitstudie zu Risikofaktoren und Diagnose chronischer Erkrankungen’’; e.g. [5]) pursues latter approach. Since the early 1990s, this epidemiological study researches personality, health, lifestyle, and cognitive variables in a population-based sample of 5 133 individuals from the German city of Heidelberg and surroundings. Based on an array of questionnaires completed at baseline that assessed depressive symptoms, resilience factors, as well as some broad personality factors (Extraversion, Neuroticism, and Psychoticism), five personality dimensions, named “The Heidelberg Five”, were subsequently extracted using exploratory factor analysis [6].

Emotional Lability (ELAB) was defined by Neuroticism, depression, low social support, low optimism, and a low sense of coherence, Lack of Behavioral Control (LBCN) by low social desirability and low anger control, Type A Behavior (TYAB) by high time urgency, exaggerated social control and Extraversion, Locus of Control over Disease (LOCC) by a high internal locus of control, and Psychoticism (PSYC) by high psychoticism (for details see [6]). On the phenotype level, ELAB is clearly associated with psychopathology (e.g. [7]). The dimension LBCN encompasses behaviors associated with executive function during development (e.g. [8]), high expression of which resemble some clinical conditions of the prefrontal cortex (e.g. [9, 10]). The personality trait TYAB [11] has been initially identified as associated with CVD, but this association was later found to have little empirical support (for a review see [12]). LOCC is a construct based on Rotter’s influential social learning theory that assesses cognitions of control over health (e.g. [13]). Importantly, external LOCC has been linked to unfavorable health behavior (e.g. [14]). PSYC is one of the three personality factors in Eysenck’s influential theory-based model of personality [15], and is being discussed as a core element of maladaptive personality, resembling schizotypy [16–18]. Recently, a subset of HeiDE participants was genotyped on whole-genome arrays, and here, we examine genomic underpinnings of The Heidelberg Five. To establish genetic similarities to and differences from the well-established Big Five trait Neuroticism, we also examined associations of The Heidelberg Five with polygenic risk scores (PRS) for Neuroticism. Also, we research longitudinal associations of The Heidelberg Five with psychopathological symptoms about 20 years after their assessment.

## METHODS AND MATERIALS

Data were analyzed using R (v3.1 or higher; [19]), PLINK 1.9 (GWAS and calculation of PRS; [20]), METAL (released 2011-03-25, meta-analyses; [21]), SHAPEIT/IMPUTE2 (imputation; [22, 23]), MAGMA (v1.07; gene, gene-set, and tissue expression analyses; [24]), and GCTA (v1.92.1beta6, estimation of SNP-based heritabilities and genetic correlations; [25]).

The analyses are covered by an ethics vote of the Medical Faculty of the University of Heidelberg (# 026/2001).

### Heidelberg Cohort Study of the Elderly

The HeiDE study is a population-based longitudinal cohort study of the inhabitants of Heidelberg (Germany) and was designed to prospectively research the association of personality and somatic diseases. Details on the baseline sample, assessed from 1992 to 1994, can be found in [6]. The final baseline sample consisted of 5 114 individuals (52.2% female) aged between 28 and 74 (99.6% between 40 and 68). Data analyzed in this study are from the baseline assessment (personality phenotypes; see below), the first follow-up (on average 8.5 years later; collection of biomaterials), and from a follow-up conducted in 2013 (psychiatric phenotypes).

### Personality assessment, principal components factor analysis and generation of factor scores

At baseline, participants completed an array of personality and health-related questionnaires (see SI). We used the original dataset of [6] and re-analyzed it using principal components followed by varimax rotation using the R *psych* library, obtaining a virtually identical solution. Regression factor scores were calculated for each latent personality dimension.

### Genotyping and imputation

DNA from saliva collected with mouthwash samples was extracted on a chemagic platform (PerkinElmer chemagen Technologie GmbH, Germany). DNA collected with Oragene OG500 Kits (DNA Genotek Inc., Canada) was extracted using DNA Genotek’s prepIT kit (DNA Genotek Inc., Canada). Samples were genotyped using two different Illumina microarrays (Illumina, San Diego, CA, USA). One subsample (HeiDE_1_) was genotyped using the Infinium PsychArray-24 BeadChip (n=2 734) and another one using the InfiniumOmniExpressExome-8v1-3_A BeadChip (HeiDE_2_; n=1 000). The combined dataset (n=3 734 pre-QC) was imputed to the 1000 Genomes phase 3 reference panel. Details on quality control (QC) and imputation can be found in the SI.

### Descriptive statistics of the genotyped sample

Of 3 320 genotyped HeiDE participants (post-QC), 34 had missing personality phenotypes, and 18 were excluded because the phenotypic sex at baseline was either missing or did not match the sex recorded at follow-up. Thus, 3 268 genotyped (HeiDE_1_: n=2 387, HeiDE_2_: n=881) were contained in the final sample. At baseline, these individuals were 52.8±7.0 (mean±SD) years old (range 28-70; 99.6% were between 40 and 68 years old), 52.3% of them were female.

### Genome-wide association studies

We conducted a GWAS for each personality phenotype. The subsamples genotyped in different batches (see above) were analyzed separately and subsequently combined using meta-analysis. The covariates for each phenotype were the following: age, sex, and the first four multidimensional scaling (MDS) components of the pairwise identity-by-state distance matrix calculated on the non-imputed genotype data.

### Gene-set and gene-property analyses

MAGMA gene-set and gene-property tissue-specific expression analysis (GTEx v7, 53 tissue types) were performed as part of the FUMA ([26]) pipeline.

### SNP-based heritabilities and genetic correlation

For each of The Heidelberg Five personality traits, we estimated the aggregate proportion of variance explained by the additive effects of all genetic SNPs/variants and genetic correlations between pairwise combinations of personality traits using GCTA GREML. We estimated the genetic relationships among all HeiDE participants, excluding cryptically related individuals with genetic similarity pi-hat>0.025, and using the same covariates as in the GWAS analyses. We used the –grm-adj 0 flag and thus assumed that causal loci have a similar distribution of allele frequencies as the genotyped SNPs.

### Calculation of polygenic risk scores

We used summary statistics of a large GWAS on Neuroticism [27] by the Social Science Genetic Association Consortium (n=170,911) as training data. PRS were calculated as the sum of the imputation dosage for each risk allele multiplied by the effect size of each genetic variant. SNPs overlapping between the Neuroticism GWAS and the HeiDE sample were clumped with an LD threshold of 0.2 within a 500 kb window. Subsequently, PRS were calculated at twelve different *p*-value thresholds (from 1×10^−6^ to 1). For each of The Heidelberg Five personality traits, we first evaluated a baseline linear regression model that predicted the factor scores of each individual personality dimension by age, sex, the first four MDS components, and the genotyping batch. We subsequently regressed the residuals of the latter model onto the Neuroticism PRS.

### The Heidelberg Five and psychiatric phenotypes at follow-up

We evaluated whether The Heidelberg Five, assessed at baseline, were associated with current depressive symptoms and lifetime anxiety phenotypes about 20 years later. The HeiDE subsample used in these analyses consisted of n=2 888 individuals, were 71.5±6.6 (mean±SD, approximated by year of birth) years old (range 53-87), and 47.4% were female (n=2 718 and n=2 660 individuals without missing data were used for analyses of depressive and anxiety symptoms, respectively). Current (past three months) depressive symptoms were assessed using a well-established 15-item questionnaire (“Allgemeine Depressionsskala”, [28], range of sum scores: 15-60). Using linear regression, we evaluated whether current depressive symptoms were associated to year of birth, sex, and factor scores of each of The Heidelberg Five measured at baseline. Visual inspection of the residuals indicted that these were not normally distributed (data not shown). We therefore log-transformed depression sum scores and subsequent visual inspection of the residuals of this model did not show obvious deviation from normality (see SI). We also tested, using logistic regression, whether a positive answer to at least one of six yes/no screening questions for lifetime anxiety symptoms (see SI) at the second follow-up were associated with year of birth, sex, and the factor scores of each of The Heidelberg Five measured at baseline. The *R*^*2*^ of both models was calculated using the R *rsq* package. For anxiety symptoms, we used a variance-function-based *R*^*2*^ for generalized linear models [29].

### Correction procedures for multiple testing

When analyzing each of The Heidelberg Five personality dimensions separately by GWAS, gene-based, gene-set and gene-property analyses, we used the conservative Bonferroni threshold to correct *p*-values, to minimize false-positives. In the analyses that compared PRS across different *p*-value thresholds, and in the analyses in which SNP-based heritabilities were compared across all personality dimensions, we used the more powerful false-discovery rate (FDR; [30]). The latter method was also used when adjusting the *p*-values of the longitudinal associations of depressive and anxiety symptoms, due to the inherent dependency of both phenotypes.

## RESULTS

### The Heidelberg Five personality dimensions

We extracted the five personality dimensions ELAB, LBCN, TYAB, LOCC, and PSYC (Figure SI1). These explained 22, 14, 10, 8, and 7% of the total variance (cumulative variance explained: 61%). The resulting factor scores had the following ranges: ELAB: -2.78 to 5.07; LBCN: -3.08 to 3.61; TYAB: -3.01 to 4.69; LOCC: -4.12 to 3.66; PSYC: -2.19 to 8.56.

### Genomic underpinnings of The Heidelberg Five

Tables 1 and 2 detail the results of SNP-based heritability analyses of and genetic correlation analyses between The Heidelberg Five. Nominally significant negative genetic correlations were found between ELAB and LBCN (Table 2) and ELAB and PSYC (Table 3), but these did not remain significant after correction for multiple testing.

**Table 1.** SNP-based heritabilities of The Heidelberg Five personality dimensions (n=2 948 for each phenotype). Abbreviations: h^2^_SNP_ – SNP-based heritability, *p*FDR – FDR-corrected *p*-value, SE – standard error.

| Phenotype | $h^2_{\text{SNP}}$ | SE | Nominal $p$ | $p_{\text{FDR}}$ |
| --- | --- | --- | --- | --- |
| ELAB | 0.339 | 0.131 | 0.004 | 0.019 |
| LBCN | 0.294 | 0.134 | 0.014 | 0.034 |
| TYAB | 0.063 | 0.132 | 0.320 | 0.320 |
| LOCC | 0.093 | 0.128 | 0.229 | 0.320 |
| PSYC | 0.079 | 0.131 | 0.275 | 0.320 |

**Table 2.** Bivariate genetic correlations between The Heidelberg Five personality dimensions (one-tailed test, n=5 896 for each phenotype pair). Abbreviations: r_G_ - genetic correlation, SE - standard error. None of the correlations survived FDR correction.

| Phenotype 1 | Phenotype 2 | $r_G$ | SE | Nominal $p$ | $p_{\text{FDR}}$ |
| --- | --- | --- | --- | --- | --- |
| ELAB | LBCN | -0.497458 | 0.345177 | 0.048914 | 0.1968200 |
| ELAB | LOCC | -0.900874 | 1.015109 | 0.059046 | 0.1968200 |
| ELAB | PSYC | -1.000000 | 0.941861 | 0.044253 | 0.1968200 |
| ELAB | TYAB | -0.700077 | 0.982098 | 0.16427 | 0.3285400 |
| LBCN | LOCC | -0.400077 | 0.730623 | 0.27359 | 0.3908429 |
| LBCN | PSYC | -0.677328 | 0.846693 | 0.15659 | 0.3285400 |
| LBCN | TYAB | 0.808054 | 1.340421 | 0.24578 | 0.3908429 |
| TYAB | LOCC | -1.000000 | 2.679146 | 0.5 | 0.5000000 |
| TYAB | PSYC | 1.000000 | 2.095205 | 0.5 | 0.5000000 |
| LOCC | PSYC | -1.0000 | 1.716760 | 0.5 | 0.5000000 |

**Table 3.** Top ten SNPs of the GWAS of PSYC. Abbreviations: Chr-chromosome, Direction-summary of effect direction for each HeiDE sample, with one ‘+’ or ‘-’ per sample, Effect-effect size for allele 1, Freq1-weighted average of frequency for allele 1 across all studies, FreqSE-corresponding standard error for allele frequency estimate, SE-overall standard error for effect size estimate, *p*-value-meta-analysis *p*-value.

| MarkerName | Chr | Position | Allele1 | Allele2 | Freq1 | FreqSE | Effect | SE | $p$-value | Direction |
| --- | --- | --- | --- | --- | --- | --- | --- | --- | --- | --- |
| rs138223660 | 8 | 130801535 | t | c | 0.986 | 0.0014 | -0.6708 | 0.1094 | 8.664e-10 | -- |
| rs112196460 | 8 | 130842384 | t | c | 0.0132 | 0.0012 | 0.6703 | 0.1098 | 1.026e-09 | ++ |
| rs113875761 | 8 | 130787390 | c | g | 0.9857 | 0.0015 | -0.6603 | 0.1089 | 1.317e-09 | -- |
| rs117161072 | 8 | 131417500 | a | g | 0.0132 | 9e-04 | 0.674 | 0.1114 | 1.459e-09 | ++ |
| rs142975048 | 8 | 130743235 | a | g | 0.9847 | 0.0014 | -0.649 | 0.1088 | 2.405e-09 | -- |
| rs147237681 | 8 | 131027478 | a | c | 0.0138 | 4e-04 | 0.6194 | 0.1051 | 3.772e-09 | ++ |
| rs139795768 | 8 | 131314036 | a | t | 0.0123 | 0.001 | 0.6684 | 0.1176 | 1.305e-08 | ++ |
| rs9882438 | 3 | 36776413 | a | g | 0.0225 | 5e-04 | 0.4953 | 0.0882 | 1.929e-08 | ++ |
| rs187876956 | 8 | 130953808 | t | c | 0.0127 | 8e-04 | 0.5997 | 0.1088 | 3.509e-08 | ++ |
| rs78247265 | 1 | 60887707 | t | c | 0.0106 | 0 | 0.8115 | 0.1523 | 9.836e-08 | +? |

### Emotional Lability

The GWAS of ELAB did not yield a genome-wide significant result (for details see the SI). Gene-based tests identified the gene *Integrin Subunit Beta 5* (*ITGB5*) as significantly associated (z=4.63, *p*=1.83×10^−6^, n=3 268), and two SNPs (rs12487905 and rs71625774) mapping to the *ITGB5* gene were among the top ten SNPs in the GWAS (both *p*<1×10^−6^, see Figure 1). Tissue expression and gene-set analyses did not yield significant results (for details see SI). The SNP-based heritability was significant (33.9%, Table 1).

**Figure 1.**
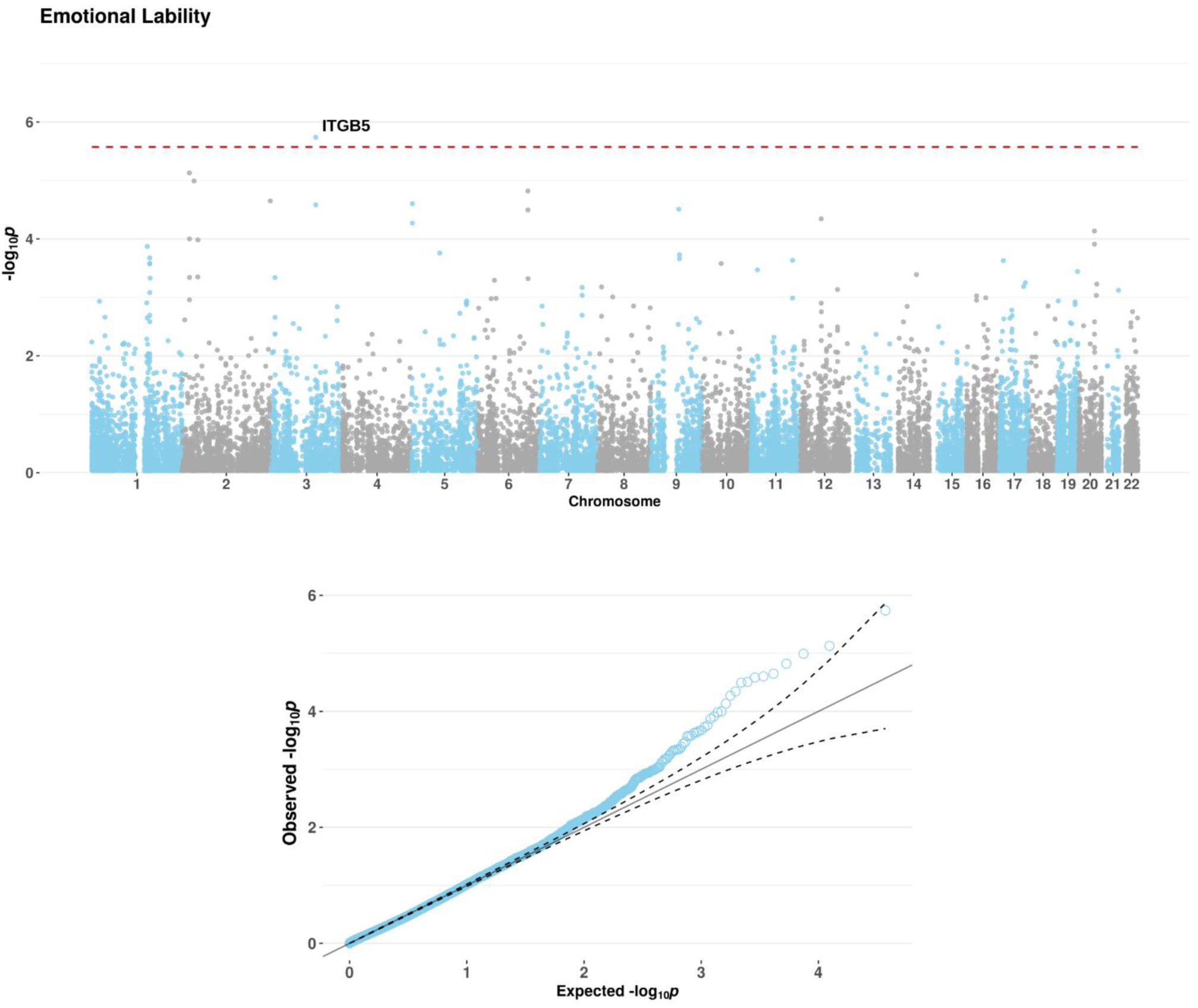
Manhattan (top) and Q-Q plots (bottom) of the gene-based test of the phenotype ELAB. Genome-wide significance level (Bonferroni-corrected for 18,634 genes) is indicated by the red dashed line.

### Low Behavioral Control

GWAS, gene-based tests, tissue expression, and gene-set analyses did not show significant results (for details see SI). We observed, however, a significant SNP-based heritability (29.4%, Table 1). Apart from the nominally significant genetic correlation with ELAB mentioned above, none of the genetic correlations between LBCN and the other personality dimensions were significant.

### Type-A Behavior

There was no genome-wide significant result for TYAB (for details see SI). The gene-based analysis identified the gene *Coiled-coil Domain Containing 83* (*CCDC83*) as significantly associated (z=4.55, *p*=2.68×10^−6^, n=3 268, see Figure 2) and two SNPs in the *CCDC83* gene (rs56160063 and rs35944027) were among the top ten SNPs in the GWAS (both *p*<2.4×10^−6^, see SI). Tissue expression and gene-set analyses did not yield significant results, neither did the SNP-bases heritability analysis nor analyses of genetic correlations between The Heidelberg Five personality dimensions (for details see SI).

**Figure 2.**
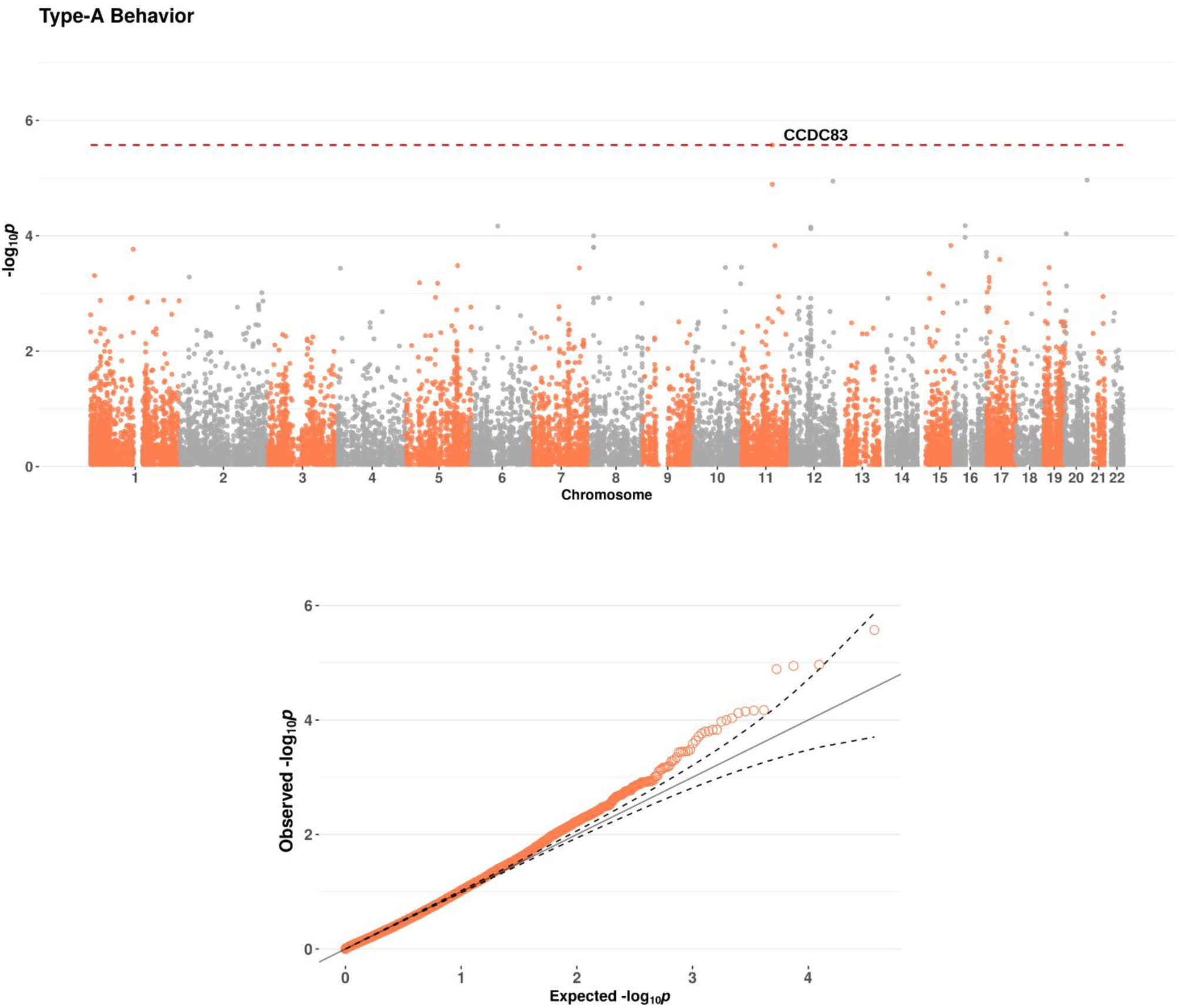
Manhattan (top) and Q-Q plots (bottom) of the gene-based test of the phenotype TYAB. Genome-wide significance level (Bonferroni-corrected for 18,634 genes) is indicated by the red dashed line.

### Locus of Control over Disease

For LOCC, GWAS, gene-based, tissue expression, SNP-based heritability, and genetic correlations with the other Heidelberg Five personality dimensions did not yield significant results (for details see SI). However, analyses identified the gene-set *GO_bp:go_cellular_response_to_ionizing_radiation* as overrepresented in the GWAS results (see SI).

### Psychoticism

The GWAS of PSYC identified a significantly associated locus on chromosome (top SNP rs138223660, *p*=8.664×10^−10^, Figure 3). Genes at this locus include *Gasdermin C* (*GSDMC*), *Family With Sequence Similarity 49 Member B* (FAM49B), and *ArfGAP With SH3 Domain, Ankyrin Repeat And PH Domain 1* (ASAP1). Another SNP on chromosome 3 (rs9882438, *p*=1.929×10^−8^), located in an intron of the *Doublecortin Like Kinase 3* (DCLK3) gene was also significantly associated with PSYC. The ten SNPs with the lowest *p*-values are shown in Table 3 (end of document). Gene-based analyses identified the gene *Nuclear Receptor Subfamily 1 Group H Member 4* (*NR1H4*, z=4.9383, *p*=3.9402×10^−7^, Figure 4) on chromosome 12 as associated with PSYC. Tissue expression, gene-set, and SNP-based heritability analyses did not find significant results.

**Figure 3.**
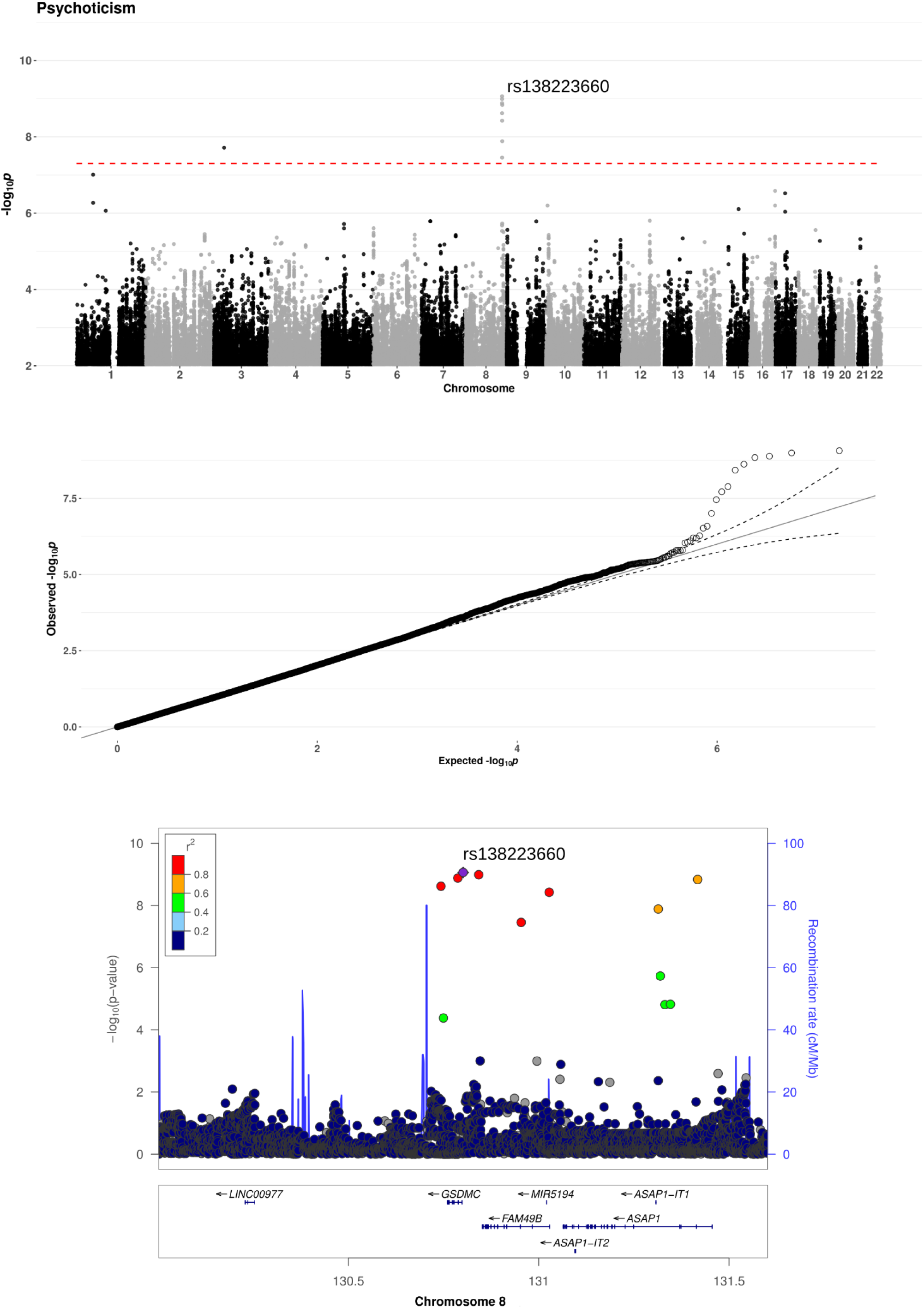
Manhattan (top), Q-Q (middle, λ= 1.007), and regional association (bottom) plots of the GWAS of the phenotype PSYC.

**Figure 4.**
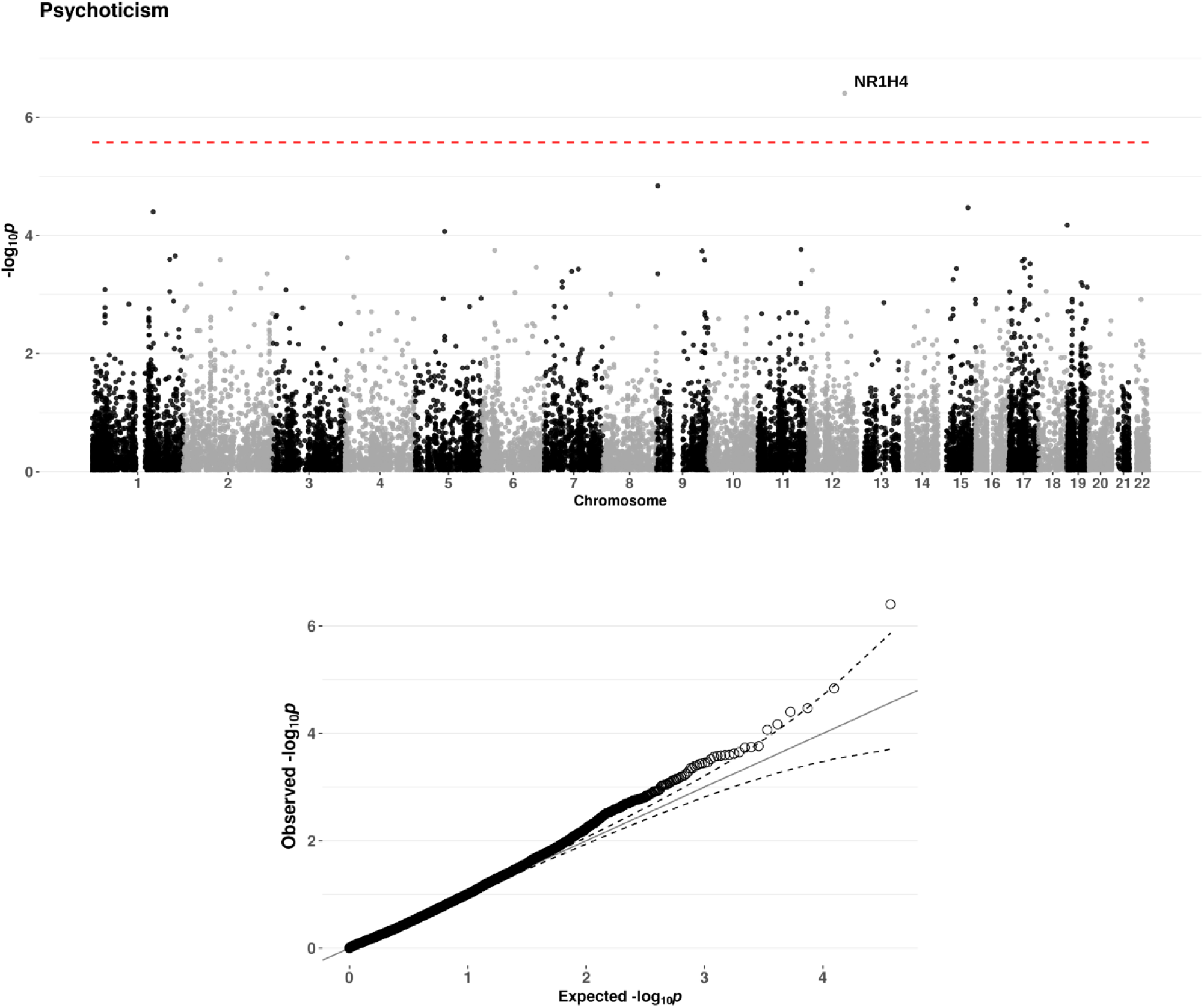
Manhattan (top) and Q-Q plots (bottom) of the gene-based test of the phenotype PSYC. Genome-wide significance level (Bonferroni-corrected for 18,634 genes) is indicated by the red dashed line.

### Associations of The Heidelberg Five with polygenic risk for neuroticism

We assessed the extent to which each of The Heidelberg Five personality dimensions shares a genetic basis with the clinically relevant Big Five personality trait Neuroticism by explaining the residuals of baseline regression models (each containing age, sex, and the first four ancestry principal components) by PRS for Neuroticism. Neuroticism PRS were significantly associated with ELAB (Figure 5), but not with the remaining personality dimensions (see SI). The direction of the association was positive, and the adjusted *R*^*2*^s of FDR-significant *p*-value thresholds (0.01, 0.05, 0.1, 0.2, 0.3, 0.4, 0.5, 1) were 0.0022, 0.0029, 0.0024, 0.0028, 0.0026, 0.0028, 0.0025, and 0.0026 (see the legend of Figure 5 for *p*-values).

**Figure 5.**
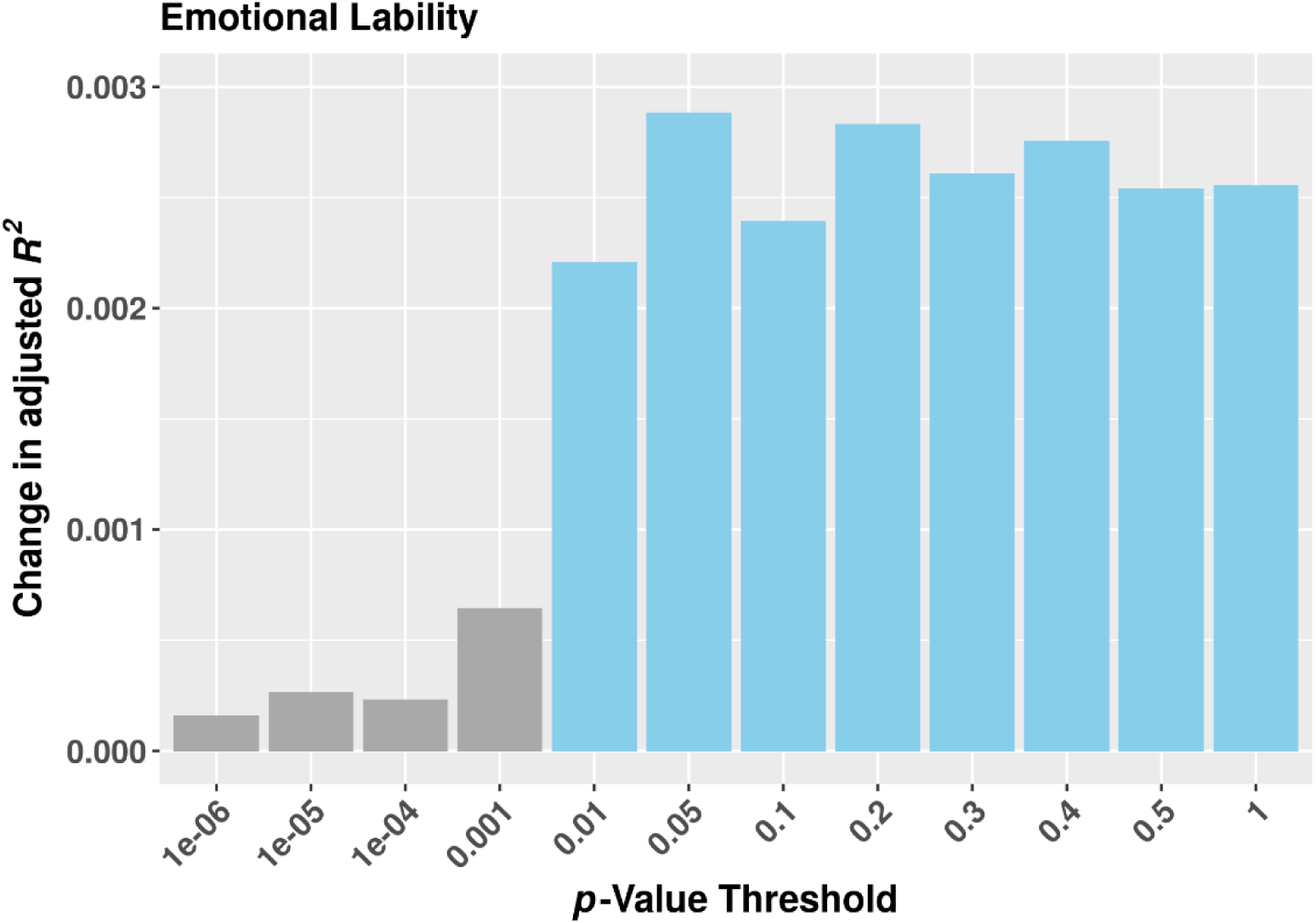
Effects (adjusted *R*^*2*^s) of PRS for neuroticism at different *p*-value thresholds on the residuals of a model regressing the personality dimension ELAB onto a set of baseline variables (see Methods and Materials). FDR-corrected *p*-values of the PRS were 0.216, 0.200, 0.200, 0.103, 0.006, 0.004, 0.005, 0.004, 0.004, 0.004, 0.004, and 0.004.

### Longitudinal associations of The Heidelberg Five and psychopathology

Table 4 (end of document) lists the result of the regression analyses. All personality dimensions showed significant longitudinal associations with current depressive symptoms about 20 years after assessment. Regarding lifetime anxiety symptoms, ELAB, LBCN, and PSYC, but not LOCC or TYAB, were significantly associated.

**Table 4.** Regression analyses of current depressive and lifetime anxiety symptoms approximately 20 years after assessment of The Heidelberg Five. Abbreviations: Std. Error - standard error of the estimate, *p*_FDR_-FDR-adjusted *p*-value. The adjusted *R*^*2*^ of the model (see text) was 31.1% for current depressive symptoms and 11.2% for lifetime anxiety symptoms. FDR-significant regressors are printed in bold font.

| | Estimate | Std. Error | t-value | $p$-value | $p_{\text{FDR}}$ |
| --- | --- | --- | --- | --- | --- |
| Intercept | 8.5428550 | 1.1990121 | 7.125 | 1.33e-12 | 5.317157e-12 |
| **Year of birth** | **-0.0027880** | **0.0006175** | **-4.515** | **6.60e-06** | **1.056070e-05** |
| **Sex (male)** | **-0.0234999** | **0.0080501** | **-2.919** | **0.00354** | **4.717726e-03** |
| **ELAB** | **0.1315498** | **0.0040296** | **32.646** | **<2e-16** | **<2e-16** |
| **LBCN** | **0.0363260** | **0.0041415** | **8.771** | **<2e-16** | **<2e-16** |
| **TYAB** | **0.0200983** | **0.0040783** | **4.928** | **8.80e-07** | **1.564546e-06** |
| **LOCC** | **-0.0084948** | **0.0040523** | **-2.096** | **0.03615** | **4.131205e-02** |
| **PSYC** | **0.0162838** | **0.0039524** | **4.120** | **3.90e-05** | **5.675753e-05** |

*Lifetime anxiety symptoms*
| | Estimate | Std. Error | z-value | $p$-value | $p_{\text{FDR}}$ |
| --- | --- | --- | --- | --- | --- |
| Intercept | -64.069467 | 12.953826 | -4.946 | 7.58e-07 | 1.556365e-06 |
| **Year of birth** | **0.032959** | **0.006671** | **4.941** | **7.78e-07** | **1.556365e-06** |
| **Sex (male)** | **-0.450115** | **0.085660** | **-5.255** | **1.48e-07** | **3.954736e-07** |
| **ELAB** | **0.613520** | **0.045513** | **13.480** | **<2e-16** | **<2e-16** |
| **LBCN** | **0.295001** | **0.044516** | **6.627** | **3.43e-11** | **1.097851e-10** |
| TYAB | 0.047679 | 0.042997 | 1.109 | 0.2675 | 2.853136e-01 |
| LOCC | 0.006232 | 0.042799 | 0.146 | 0.8842 | 8.842379e-01 |
| **PSYC** | **0.110470** | **0.043120** | **2.562** | **0.0104** | **1.281183e-02** |

## DISCUSSION

The aim of this work was to further characterize The Heidelberg Five using information on common genetic variants and long-term follow-up data, to gain a more comprehensive understanding of both their biological basis and their putative importance in predicting longitudinal outcomes. Regarding the follow-up analysis, it was surprising that each of The Heidelberg Five (high ELAB, low behavioral control, high TYAB, low internal LOCC and high PSYC) was associated with more severe depressive symptoms, measured at the 20-year follow-up. These findings alone corroborate the importance of health-related personality traits, providing justification for further research. Different SNP-based heritabilities across The Heidelberg Five furthermore suggest a varying importance of common genetic variants, albeit this may depend on the population under study [31]. Specifically, both ELAB and LBCN show substantial SNP-based heritabilities, as do the Big Five traits Neuroticism (*h*^*2*^_SNP_ =15%) and Openness to Experience (*h*^*2*^_SNP_ =21%, [32]). In the latter study, only the two The Big Five traits mentioned showed substantial h^2^_SNP_, and, in the present study, we did not detect significant h^2^_SNP_ for TYAB, LOCC, or PSYC. Similar to [32], we may interpret this as emphasizing the putative importance of rare or structural variants for these personality dimensions, as all personality phenotypes are heritable to some degree [33]. Also, it is possible that unknown environmental covariates exist that explain more phenotypic variance of TYAB, LOCC, and PSYC, and accounting for these would result in larger observed SNP-based heritabilities also for these personality dimensions.

The orthogonality of The Heidelberg Five on the phenotype level is reflected by non-significant genetic correlations between them. Conversely, both the phenotypic and genotypic relatedness of Neuroticism and ELAB is reflected in substantial SNP-based heritabilities of both traits (see above) and by the result that Neuroticism PRS explain variation of ELAB. This was not the case for the remaining Heidelberg Five personality dimensions. We further discuss the results of biological analyses of each personality dimension below.

### Emotional Lability

Both GWAS and gene-based analyses identified *ITGB5*, encoding a transmembrane protein. The family of integrins, to which *ITGB5* belongs, are membrane proteins that translate intracellular signaling to extracellular interactions. They have been associated with neuropsychiatric disease [34] and coordinate both synaptic structure and function [35]. Furthermore, SNPs in the *ITGB5* gene are associated with blood pressure [36] and coronary artery disease [37]. Interestingly, an association between ELAB and the incidence of CVD was previously identified in longitudinal analyses ([6]) and thus may suggest a common genetic basis of both. Concerning the longitudinal associations of high ELAB scores with both depressive and anxiety symptoms, observed in the present study, confirm the well-known clinical importance of this Neuroticism-like phenotype [38], and are in line with meta-analyses of longitudinal studies of Neuroticism [39, 40].

### Lack of Behavioral Control

LBCN is a latent personality trait that is negatively correlated with anger control and social desirability and positively correlated with aggression, irritability, and outwards expression of anger [6]. These personality facets are of critical importance for self-regulation, an intensively studied construct in the behavioral sciences. Specifically, effective self-regulation allows individuals to plan and to focus beyond immediate needs, relying heavily on proper functioning of the prefrontal cortex. As such, LBCN may be regarded as an “executive function” personality trait. Indeed, better performance on executive tasks, such as the Wisconsin-Card-Sorting or the Stroop Test, is associated with higher scores on the Big Five dimensions Agreeableness and Conscientiousness [41], which describe phenotypically similar higher-order personality constructs in the Big Five framework. As mentioned above, LBCN showed a relatively high *h*^*2*^_SNP_ which is supported by previous twin studies that found differences in specific executive control functions to be almost entirely genetic in origin [42].

### Type-A Behavior

Psychological tests measuring time urgency, exaggerated social control, and extraversion (for details see SI) loaded highly on the personality dimension TYAB. Type-A behavior is “characterized primarily by a chronic incessant struggle to achieve more and more in less and less time” ([43]), and was initially hypothesized to be associated with CVD, but this assumption was later found to have little empirical support. In line with this finding, prospective research in the HeiDE study did not identify TYAB as a predictor of the incidence of CVD [6]. Regarding biological correlates, gene-based analysis pointed to the protein-coding gene coiled-coil domain containing 83 (CCDC83). In European populations, this gene has been linked to urinary tract infection frequency [44], but not to behavioral phenotypes.

### Locus of Control over Disease

LOCC is a personality dimension derived from Rotter’s influential social learning theory (e.g. [45]). Briefly, it can be shown that individuals differ in their perception of reinforcements and classify these either as being controlled externally, i.e., by chance or the specific situation, or internally, by the person’s own actions. According to Rotter, generalized expectancies differ between individuals and this constitutes a personality dimension, Locus of Control. This personality dimension determines whether individuals perceive outcomes as rather externally or internally controlled [46]. High internal Locus of Control has been shown to be important for a variety of health behaviors including smoking, alcohol consumption, exercise, diet [47, 48], and medication adherence [14]. GWAS and gene-based analyses did not detect significant results. This is in line with Rotter’s view that Locus of Control is a consequence of learning experiences. Consistently, the construct also exhibits a malleable, state-like property [49]. However, the gene-set *GO_bp:go_cellular_response_to_ionizing_radiation* was overrepresented amongst the LOCC results (see SI). Among the 51 genes in this gene-set, five are classified as transcription factors, one as cell differentiation marker, four as protein kinases, and four as tumor suppressors in MSigDB [50]. Transcription factors have previously been linked to health-relevant personality traits [51, 52], and are crucial for long-term memory formation [53]. Thus, LOCC may be characterized by transcriptional alterations that mediate the attribution of outcomes as being either externally or internally controlled.

### Psychoticism

Alongside Extraversion and Neuroticism, Psychoticism constitutes the third factor of Eysenck’s P-E-N model of personality that is based on biological and experimental grounds [18]. Initially, Psychoticism was conceptualized as a continuous dimension, predisposing individuals to psychosis, but a 10-year longitudinal study did not confirm this association with psychosis [16]. The latter study, however, also reported that individuals scoring high on Psychoticism “exceeded controls on ratings of psychotic-like experiences and on symptoms of schizotypal and paranoid personality disorder”. Furthermore, based on a number of analyses, Zuckerman described the personality dimension Psychoticism as encompassing “impulsivity, lack of socialization and responsibility, aggression, a strong need for independence, and sensation seeking”, with clinical extremes [54]. While no significant SNP-based heritability of this trait was detected, GWAS and gene-based analysis revealed significant loci on chromosomes 3, 8, and 12. Of the genes in these loci, a SNP in *FAM49B* showed a suggestive association with post-traumatic stress disorder [55]. Furthermore, SNPs in ASAP1 were suggestively associated with autism spectrum disorder [56] and, in individuals with Ashkenazi Jewish ancestry, suggestively associated with schizophrenia [57].

Several findings emerge from the present analyses of The Heidelberg Five. Firstly, each personality dimension is associated with psychiatric phenotypes, measured some 20 years later, which underlines their clinical significance. Secondly, ELAB is genetically related Neuroticism. As ELAB explained most of the phenotypic variance in the factor analysis, the behavioral importance of a Neuroticism/ELAB phenotype is further underscored. Thirdly, LBCN, a previously unknown latent “executive function” personality dimension has emerged as a heritable trait, warranting further investigation.

Our results need to be interpreted keeping several limitations in mind. While based on longitudinal data, we used cross-sectional analyses ignoring accrual and mortality. If any of the traits or SNPs are associated with accrual or mortality, this will introduce selection bias. Results of the effects of psychological traits on CVD and cancer, including cause-specific mortality, are reassuring, however, insofar as most had no major impact on these outcomes [5].

Supplementary information is available.

## ACKNOWLEDGEMENTS

Thomas G. Schulze is supported by the Deutsche Forschungsgemeinschaft (DFG) within the framework of the projects www.kfo241.de and www.PsyCourse.de (SCHU 1603/4-1, 5-1, 7-1; FA241/16-1) and by the German Federal Ministry of Education and Research (BMBF) through the Integrated Network IntegraMent (Integrated Understanding of Causes and Mechanisms in Mental Disorders), under the auspices of the e:Med Programme with grants awarded to Thomas G. Schulze (01ZX1614K), Marcella Rietschel (01ZX1614G), Markus M. Nöthen (01ZX1614A), and Bertram Müller-Myhsok (01ZX1614J). Marcella Rietschel is additionally supported by the ERA-NET NEURON grants “SynSchiz” (01EW1810) and “EMBED” (01EW1904). Thomas G. Schulze received additional support from the German Federal Ministry of Education and Research (BMBF) within the framework of the BipoLife network and the Dr. Lisa Oehler Foundation, Kassel (Germany). Franziska Degenhardt received support from the BONFOR Programme of the University of Bonn, Germany.

## REFERENCES

1. Sanchez-Roige S, Gray JC, MacKillop J, Chen C-H, Palmer AA. The genetics of human personality. Genes Brain Behav. 2018;17:e12439.

2. Goldberg LR. The structure of phenotypic personality traits. Am Psychol. 1993;48:26–34.

3. Capitanio JP. Personality and disease. Brain Behav Immun. 2008;22:647–650.

4. Friedman HS, Booth-Kewley S. Personality, type A behavior, and coronary heart disease: the role of emotional expression. J Pers Soc Psychol. 1987;53:783–792.

5. Stürmer T, Hasselbach P, Amelang M. Personality, lifestyle, and risk of cardiovascular disease and cancer: follow-up of population based cohort. BMJ. 2006;332:1359.

6. Amelang M, Hasselbach P, Stürmer T. Personality, Cardiovascular Disease, and Cancer: First Results from the Heidelberg Cohort Study of the Elderly. Z Gesundheitspsychol; 12:102–115.

7. Ormel J, Jeronimus BF, Kotov R, Riese H, Bos EH, Hankin B, et al. Neuroticism and common mental disorders: meaning and utility of a complex relationship. Clin Psychol Rev. 2013;33:686–697.

8. Rohlf HL, Holl AK, Kirsch F, Krahé B, Elsner B. Longitudinal Links between Executive Function, Anger, and Aggression in Middle Childhood. Front Behav Neurosci. 2018;12:27.

9. Szczepanski SM, Knight RT. Insights into human behavior from lesions to the prefrontal cortex. Neuron. 2014;83:1002–1018.

10. Tate RL. Executive dysfunction and characterological changes after traumatic brain injury: two sides of the same coin? Cortex. 1999;35:39–55.

11. Friedman M, Rosenman RH. Association of specific overt behavior pattern with blood and cardiovascular findings; blood cholesterol level, blood clotting time, incidence of arcus senilis, and clinical coronary artery disease. J Am Med Assoc. 1959;169:1286–1296.

12. Kuper H, Marmot M, Hemingway H. Systematic review of prospective cohort studies of psychosocial factors in the etiology and prognosis of coronary heart disease. Semin Vasc Med. 2002;2:267–314.

13. Wooldridge KL, Wallston KA, Graber AL, Brown AW, Davidson P. The relationship between health beliefs, adherence, and metabolic control of diabetes. Diabetes Educ. 1992;18:495–500.

14. Náfrádi L, Nakamoto K, Schulz PJ. Is patient empowerment the key to promote adherence? A systematic review of the relationship between self-efficacy, health locus of control and medication adherence. PLoS ONE. 2017;12:e0186458.

15. Eysenck HJ, Eysenck SBG. Psychoticism as a dimension of personality. London: Hodder and Stoughton; 1976.

16. Chapman JP, Chapman LJ, Kwapil TR. Does the Eysenck psychoticism scale predict psychosis? A ten year longitudinal study. Personality and Individual Differences. 1994;17:369–375.

17. Wright AGC, Thomas KM, Hopwood CJ, Markon KE, Pincus AL, Krueger RF. The hierarchical structure of DSM-5 pathological personality traits. J Abnorm Psychol. 2012;121:951–957.

18. van Kampen D. Personality and Psychopathology: a Theory-Based Revision of Eysenck’s PEN Model. Clin Pract Epidemiol Ment Health. 2009;5:9–21.

19. R Core Team. R: A Language and Environment for Statistical Computing. Vienna, Austria: R Foundation for Statistical Computing; 2014.

20. Chang CC, Chow CC, Tellier LC, Vattikuti S, Purcell SM, Lee JJ. Second-generation PLINK: rising to the challenge of larger and richer datasets. Gigascience. 2015;4:7.

21. Willer CJ, Li Y, Abecasis GR. METAL: fast and efficient meta-analysis of genomewide association scans. Bioinformatics. 2010;26:2190–2191.

22. Howie BN, Donnelly P, Marchini J. A flexible and accurate genotype imputation method for the next generation of genome-wide association studies. PLoS Genet. 2009;5:e1000529.

23. Delaneau O, Marchini J, Zagury J-F. A linear complexity phasing method for thousands of genomes. Nat Methods. 2012;9:179–181.

24. de Leeuw CA, Mooij JM, Heskes T, Posthuma D. MAGMA: generalized gene-set analysis of GWAS data. PLoS Comput Biol. 2015;11:e1004219.

25. Yang J, Lee SH, Goddard ME, Visscher PM. GCTA: a tool for genome-wide complex trait analysis. Am J Hum Genet. 2011;88:76–82.

26. Watanabe K, Taskesen E, van Bochoven A, Posthuma D. Functional mapping and annotation of genetic associations with FUMA. Nat Commun. 2017;8:1826.

27. Okbay A, Baselmans BML, De Neve J-E, Turley P, Nivard MG, Fontana MA, et al. Genetic variants associated with subjective well-being, depressive symptoms, and neuroticism identified through genome-wide analyses. Nat Genet. 2016;48:624–633.

28. Hautzinger M, Bailer M. Allgemeine Depressions Skala: ADS. Weinheim: Beltz; 1993.

29. Zhang D. A Coefficient of Determination for Generalized Linear Models. The American Statistician. 2017;71:310–316.

30. Benjamini Y, Hochberg Y. Controlling the False Discovery Rate: A Practical and Powerful Approach to Multiple Testing. Journal of the Royal Statistical Society: Series B (Methodological). 1995;57:289–300.

31. Moore DS, Shenk D. The heritability fallacy. Wiley Interdiscip Rev Cogn Sci. 2017;8.

32. Power RA, Pluess M. Heritability estimates of the Big Five personality traits based on common genetic variants. Transl Psychiatry. 2015;5:e604.

33. Turkheimer E, Pettersson E, Horn EE. A phenotypic null hypothesis for the genetics of personality. Annu Rev Psychol. 2014;65:515–540.

34. Carneiro AMD. The emerging role of integrins in neuropsychiatric disorders. Neuropsychopharmacology. 2010;35:338–339.

35. Park YK, Goda Y. Integrins in synapse regulation. Nat Rev Neurosci. 2016;17:745–756.

36. Giri A, Hellwege JN, Keaton JM, Park J, Qiu C, Warren HR, et al. Trans-ethnic association study of blood pressure determinants in over 750,000 individuals. Nat Genet. 2019;51:51–62.

37. Nelson CP, Goel A, Butterworth AS, Kanoni S, Webb TR, Marouli E, et al. Association analyses based on false discovery rate implicate new loci for coronary artery disease. Nat Genet. 2017;49:1385–1391.

38. Gale CR, Hagenaars SP, Davies G, Hill WD, Liewald DCM, Cullen B, et al. Pleiotropy between neuroticism and physical and mental health: findings from 108 038 men and women in UK Biobank. Transl Psychiatry. 2016;6:e791.

39. Jeronimus BF, Kotov R, Riese H, Ormel J. Neuroticism’s prospective association with mental disorders halves after adjustment for baseline symptoms and psychiatric history, but the adjusted association hardly decays with time: a meta-analysis on 59 longitudinal/prospective studies with 443 313 participants. Psychol Med. 2016;46:2883–2906.

40. Hakulinen C, Elovainio M, Pulkki-Råback L, Virtanen M, Kivimäki M, Jokela M. PERSONALITY AND DEPRESSIVE SYMPTOMS: INDIVIDUAL PARTICIPANT META-ANALYSIS OF 10 COHORT STUDIES: Research Article: Personality and Depression. Depress Anxiety. 2015;32:461–470.

41. Jensen-Campbell LA, Rosselli M, Workman KA, Santisi M, Rios JD, Bojan D. Agreeableness, conscientiousness, and effortful control processes. Journal of Research in Personality. 2002;36:476–489.

42. Friedman NP, Miyake A, Young SE, DeFries JC, Corley RP, Hewitt JK. Individual differences in executive functions are almost entirely genetic in origin. J Exp Psychol Gen. 2008;137:201–225.

43. Smith DF, Sterndorff B, Røpcke G, Gustavsen EM, Hansen JK. Prevalence and severity of anxiety, depression and Type A behaviors in angina pectoris. Scand J Psychol. 1996;37:249–258.

44. Tian C, Hromatka BS, Kiefer AK, Eriksson N, Noble SM, Tung JY, et al. Genome-wide association and HLA region fine-mapping studies identify susceptibility loci for multiple common infections. Nat Commun. 2017;8:599.

45. Rotter JB. Generalized expectancies for internal versus external control of reinforcement. Psychol Monogr. 1966;80:1–28.

46. Weiner B, Reisenzein R, Pranter W. Motivationspsychologie. 3. Aufl., [unveränd. Nachdr.]. Weinheim: Beltz, Psychologie-Verl.-Union; 2009.

47. Steptoe A, Wardle J. Locus of control and health behaviour revisited: a multivariate analysis of young adults from 18 countries. Br J Psychol. 2001;92:659–672.

48. Strudler Wallston B, Wallston KA. Locus of Control and Health: A Review of the Literature. Health Education Monographs. 1978;6:107–117.

49. Galvin BM, Randel AE, Collins BJ, Johnson RE. Changing the focus of locus (of control): A targeted review of the locus of control literature and agenda for future research. J Organ Behav. 2018;39:820–833.

50. Liberzon A, Subramanian A, Pinchback R, Thorvaldsdottir H, Tamayo P, Mesirov JP. Molecular signatures database (MSigDB) 3.0. Bioinformatics. 2011;27:1739–1740.

51. Damberg M, Garpenstrand H, Alfredsson J, Ekblom J, Forslund K, Rylander G, et al. A polymorphic region in the human transcription factor AP-2β gene is associated with specific personality traits. Mol Psychiatry. 2000;5:220–224.

52. Damberg M, Berggård C, Mattila-Evenden M, Rylander G, Forslund K, Garpenstrand H, et al. Transcription Factor AP-2β Genotype Associated with Anxiety-Related Personality Traits in Women. Neuropsychobiology. 2003;48:169–175.

53. Alberini CM. Transcription Factors in Long-Term Memory and Synaptic Plasticity. Physiological Reviews. 2009;89:121–145.

54. Zuckerman M. Personality in the third dimension: A psychobiological approach. Personality and Individual Differences. 1989;10:391–418.

55. Xie P, Kranzler HR, Yang C, Zhao H, Farrer LA, Gelernter J. Genome-wide association study identifies new susceptibility loci for posttraumatic stress disorder. Biol Psychiatry. 2013;74:656–663.

56. Grove J, Ripke S, Als TD, Mattheisen M, Walters RK, Won H, et al. Identification of common genetic risk variants for autism spectrum disorder. Nat Genet. 2019;51:431–444.

57. Goes FS, McGrath J, Avramopoulos D, Wolyniec P, Pirooznia M, Ruczinski I, et al. Genome-wide association study of schizophrenia in Ashkenazi Jews. Am J Med Genet B Neuropsychiatr Genet. 2015;168:649–659.

