## Supplemental Material for "“The Heidelberg Five” Personality Dimensions: Genome-wide Associations, Polygenic Risk for Neuroticism, and Psychopathology 20 Years after Assessment"

**Supplementary Information**

#### *1. Personality constructs and tests used at the baseline assessment*

- Time Urgency and Perpetual Activation Scale (Wright, L., McCurdy, S., & Rogoll, G. (1992). The TUPA scale: a self-report measure for the Type A subcomponent of time urgency and perpetual activation. Unpublished manuscript, University of Oklahoma, OK.)
- State-Trait-Anger Expression-Inventory (24 items measuring “Anger Out”, “Anger In” and “Anger control”, Schwenkmezger, P., Hodapp, V., & Spielberger, C. D. (1992). Das State-Trait-Ärgerausdrucks-Inventar (STAXI). Bern: Huber.)
- Hostility (Dimensions “Aggression”, “Irritability” and “Jealousy” from Kornadt, H. J. (1982). Aggressionsmotiv und Aggressionshemmung (2. Aufl.). Stuttgart: Huber., see Amelang 2004 for details)
- Exaggerated social control (Way of Life Scale, Wright, L., Bussmann, K. von, Freidman, A., Khoury, M., Owens, F., & Paris, W. (1990). Exaggerated social control and its relationship to the Type A behaviour pattern. Journal of Research in Personality, 24, 258–269.)
- Depression Scale (Zerssen, D. von (1976). Die Paranoid-Depressivitäts-Skala. Weinheim: Beltz-Test)
- Sense of Coherence Scale (Schmidt-Rathjens, C., Benz, D., Van Damme, D., Feldt, K., & Amelang, M. (1997). Über zwiespältige Erfahrungen mit Fragebögen zum Kohärenzsinn. Diagnostica, 43, 327–346.)
- Optimism (Life Orientation Test, Scheier, M. F. & Carver, C. S. (1985). Optimism, coping and health: Assessment and implications of generalised outcome expectancies. Health Psychology, 4, 219–247.)
- Questionnaire for measuring the locus of control over diseases (Ferring, D. & Filipp, S. H. (1989). Der Fragebogen zur Erfassung gesundheitsbezogener Kontrollüberzeugungen (FEGK) [Questionnaire to measure belief in control over one’s own health status]. Zeitschrift für Klinische Psychologie, 28, 285–289.)
- Social Support-Scale (Fydrich, T., Sommer, G., Menzel, G., & Höll, B. (1987). Fragebogen zur sozialen Unterstützung (Kurzform; SOZU-K-22). Zeitschrift für Klinische Psychologie, 16, 434–436.)
- Eysenck-Personality-Inventory (Extraversion, Neuroticism, and Social Desirability; Eggert, D. (1974). Eysenck-Persönlichkeits-Inventar (EPI). Göttingen: Hogrefe.)

- Psychoticism (Baumann, U. & Dittrich, A. (1976). Überprüfung der Fragebogendimension P (Psychotizismus) im Vergleich zu Extraversion und Neurotizismus. Zeitschrift für Klinische Psychologie, 5, 1–22).

***2. Screeplot of the principal components analysis that led to the identification of The Heidelberg Five***

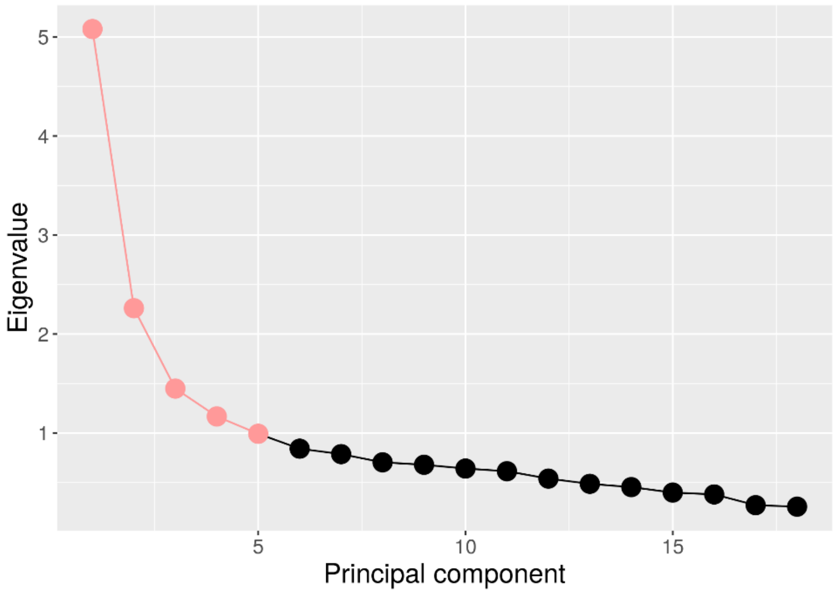


Figure SI1. Screeplot of principal components in the factor analysis from which The Heidelberg Five (red) were extracted.

#### *3. Genotype quality control (QC)*

QC of genotype data was conducted in PLINK v1.90b3.36 or higher. QC was carried out first on each of both subsamples separately (HeiDE$_1$ and HeiDE$_2$), followed by a second round of QC on the combined dataset.

*Sequence of QC steps:*

1. HeiDE1
   Before QC: 2 734 individuals and 588 454 variants
1.1. Removal of SNPs with call rates <98% or a MAF <1%
1.2. Removal of individuals with genotyping rates <98% (32 removed)
1.3. Removal of gender mismatches (42 removed)
1.4. Removal of genetic duplicates (4 removed)
1.5. Removal of cryptic relatives with pi-hat≥12.5 (112 removed)
1.6. Removal of genetic outliers with a distance from the mean of >4 SD in the first eight MDS components (26 removed)
1.7. Removal of individuals with a deviation of the autosomal or X-chromosomal heterozygosity from the mean >4 SD (15 removed)
1.8. Removal of non-autosomal variants
1.9. Removal of SNPs with call rates <98% or a MAF <1% or Hardy-Weinberg Equilibrium (HWE) test p-values <1×10-6
1.10. Removal of A/T and G/C SNPs
1.11. Update of variant IDs and positions to the IDs and positions in the 1000 Genomes Phase 3 reference panel
1.12. Alignment of alleles to the reference panel
1.13. Removal of duplicated variants and variants not present in the reference panel
   After QC: 2 503 individuals and 278 307 variants

2. HeiDE2
   Before QC: 1,000 individuals and 958 497 variants
2.1. Removal of SNPs with call rates <98% or a MAF <1%
2.2. Removal of individuals with genotyping rates <98% (6 removed)
2.3. Removal of gender mismatches (14 removed)
2.4. Removal of genetic duplicates (1 removed)
2.5. Removal of cryptic relatives with pi-hat≥12.5 (20 removed)
2.6. Removal of genetic outliers with a distance from the mean of >4 SD in the first eight MDS components (18 removed)

2.7. Removal of individuals with a deviation of the autosomal or X-chromosomal heterozygosity from the mean >4 SD (8 removed)

2.8. Removal of non-autosomal variants

2.9. Removal of SNPs with call rates <98% or a MAF <1% or HWE test p-values <1×10-6

2.10. Removal of A/T and G/C SNPs

2.11. Update of variant IDs and positions to the IDs and positions in the 1000 Genomes Phase 3 reference panel

2.12. Alignment of alleles to the reference panel

2.13. Removal of duplicated variants and variants not present in the reference panel

After QC: 933 individuals and 628 383 variants

3. Combined dataset of both samples (3 436 individuals and 251 777 variants)

3.1. Removal of SNPs with call rates <98% or a MAF <1%

3.2. Removal of individuals with genotyping rates <98% (1 removed)

3.3. Removal of genetic duplicates caused by an overlap between both datasets (44 removed)

3.4. Removal of cryptic relatives with pi-hat≥12.5 (50 removed)

3.5. Removal of genetic outliers with a distance from the mean of >4 SD in the first eight MDS components (12 removed)

3.6. Removal of individuals with a deviation of the autosomal heterozygosity from the mean >4 SD (9 removed from the FAM sample)

3.7. Removal of SNPs with call rates <98% or a MAF <1% or HWE test p-values <1×10-6

After QC: 3 320 individuals and 251 642 variants

*Imputation of genotype data*

Genotypes were aligned to the 1000 Genomes Phase 3 reference panel using SHAPEIT v2 (r837) and PLINK v1.90b3.44. Pre-phasing (haplotype estimation) was conducted for each chromosome separately using SHAPEIT. Imputation was performed using IMPUTE2 v2.3.2 in 5 Mbp chunks with 500 kb buffers, filtering out variants that are monomorphic in the EUR samples. Chunks with <51 genotyped variants or concordance rates <92 % were fused with neighboring chunks and re-imputed. Imputed variants with a MAF <1% or an INFO metric <0.8 were removed.

Imputed variants in the combined sample after QC: 8 348 464

#### *4. Results of the phenotype Emotional Lability (ELAB)*

Manhattan and Q-Q plots of SNP data, and the results of gene-property tissue-specific expression analysis (MAGMA; GTEx v6, 53 tissue types) are shown in Figures SI2-SI4. Top 10 GWAS SNPs are listed in Table SI1, Table SI2 shows the top 10 associated gene-sets from MAGMA gene-set analysis.

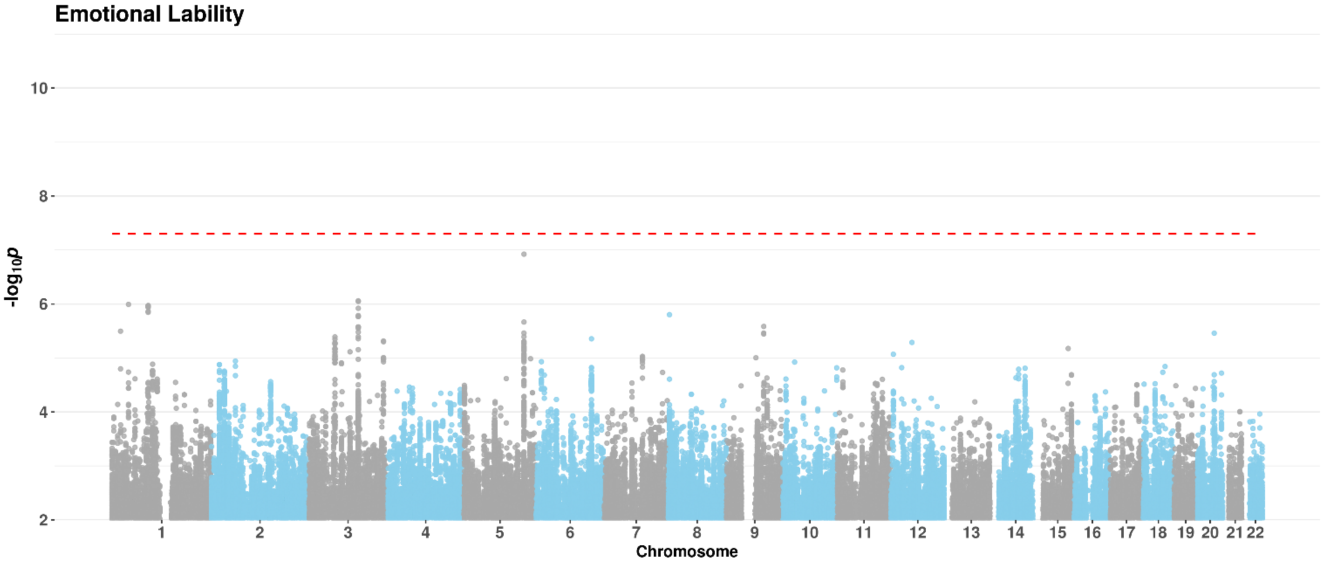


Figure SI2. Manhattan plot of SNP data of the phenotype ELAB. Genome-wide significance level is indicated by the red dashed line.

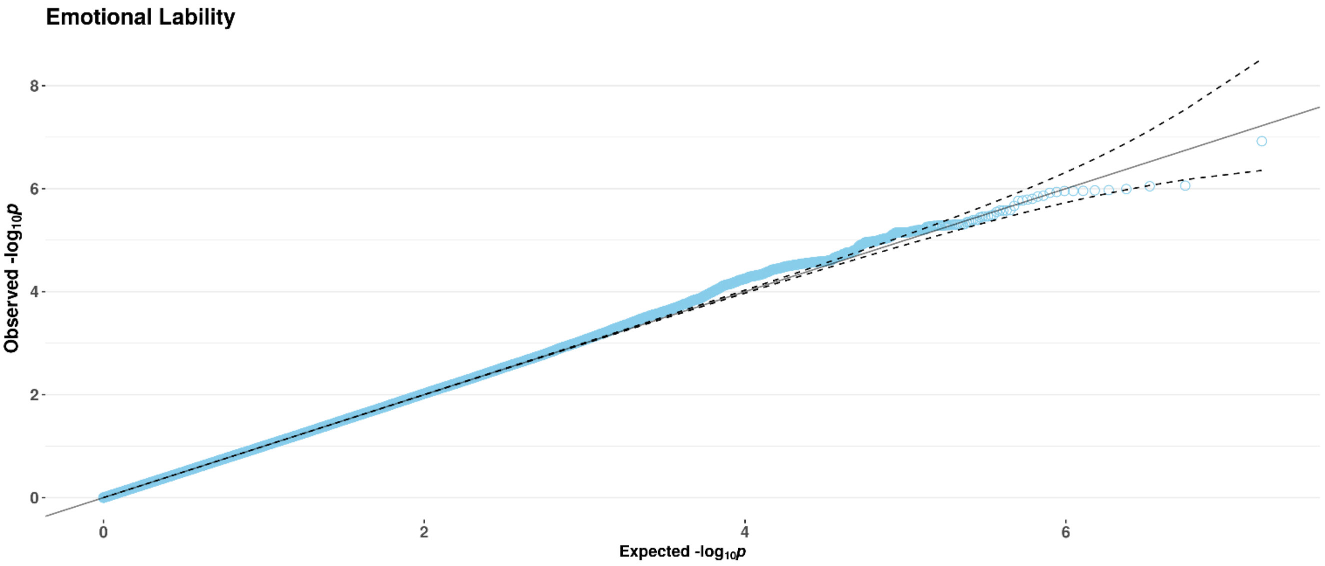


Figure SI3. Q-Q plot of SNP data of the phenotype ELAB (λ=0.997).

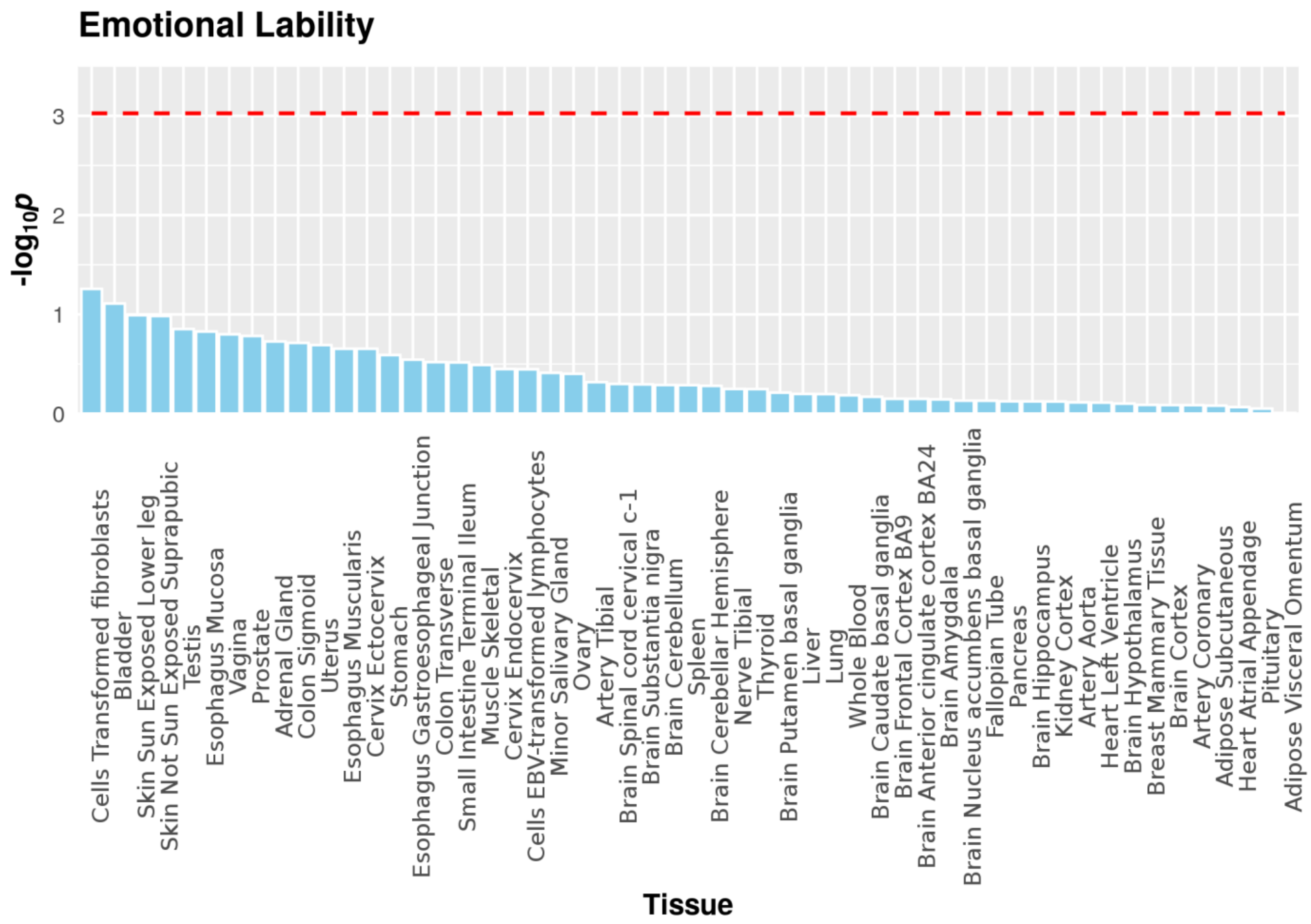


Figure SI4. Results of MAGMA gene-property tissue expression analysis of the phenotype ELAB. Bonferroni-corrected significance level (for 53 tissues) is indicated by the red dashed line.

Table SI1. Top ten SNPs from the GWAS of the phenotype ELAB (meta-analysis of both HeiDE samples). Abbreviations: Chr-chomosome, Effect-beta for allele 1, Freq1- weighted average of frequency for allele 1 across all studies, FreqSE-corresponding standard error for allele frequency estimate, SE-overall standard error for effect size estimate, *p*-value-meta-analysis *p*-value, Direction-summary of effect direction for each HeiDE sample, with one '+' or '-' per sample.

| MarkerName | Chr | Position | Allele1 | Allele2 | Freq1 | FreqSE | Effect | SE | *p*-value | Direction |
|---|---|---|---|---|---|---|---|---|---|---|
| rs2344174 | 5 | 150341653 | t | g | 0.8566 | 0.001 | -0.187 | 0.0353 | 1.196e-07 | -- |
| rs12487905 | 3 | 124521035 | t | c | 0.8019 | 0.0054 | -0.1528 | 0.0311 | 8.745e-07 | -- |
| rs71625774 | 3 | 124533325 | t | ta | 0.2408 | 0.0082 | 0.152 | 0.0309 | 8.964e-07 | ++ |
| rs34351096 | 1 | 41858649 | ct | c | 0.547 | 0.0053 | 0.1264 | 0.0259 | 1.018e-06 | ++ |
| rs17131127 | 1 | 91135628 | t | c | 0.1333 | 5e-04 | 0.1753 | 0.0359 | 1.068e-06 | ++ |
| rs112852264 | 1 | 91151681 | t | g | 0.1334 | 6e-04 | 0.1748 | 0.0358 | 1.08e-06 | ++ |
| rs34464186 | 1 | 91149374 | t | c | 0.8667 | 5e-04 | -0.1746 | 0.0358 | 1.103e-06 | -- |
| rs12746528 | 1 | 91146474 | t | c | 0.8667 | 5e-04 | -0.1746 | 0.0358 | 1.11e-06 | -- |
| rs66583506 | 1 | 91140999 | a | t | 0.1333 | 5e-04 | 0.1746 | 0.0358 | 1.11e-06 | ++ |
| rs12750662 | 1 | 91146654 | a | t | 0.1334 | 6e-04 | 0.1743 | 0.0358 | 1.158e-06 | ++ |

Table SI2. Top 10 gene-sets associated with the phenotype ELAB (meta-analysis of both HeiDE samples). Abbreviations: NGenes- the number of genes in the data that are in the gene set, Beta- regression coefficient of the gene set, Beta STD-the semi-standardized regression coefficient, corresponding to the predicted change in Z-value given a change of one standard deviation in the predictor gene set, SE-the standard error of the regression coefficient, *p*-value- the competitive gene-set p-value, $p_{Bon}$-Bonferroni-corrected p-value.

| Gene Set | NGenes | Beta | Beta STD | SE | *p*-value | $p_{Bon}$ |
|---|---|---|---|---|---|---|
| GO_bp:go_protein_acylation | 140 | 0.23239 | 0.020067 | 0.065472 | 0.00019352 | 1 |
| GO_bp:go_linoleic_acid_metabolic_process | 13 | 0.79105 | 0.020886 | 0.22669 | 0.0002425 | 1 |
| Curated_gene_sets:yamashita_liver_cancer_with_epcam_dn | 15 | 0.65781 | 0.018655 | 0.19331 | 0.00033429 | 1 |
| Curated_gene_sets:motamed_response_to_androgen_up | 5 | 0.95485 | 0.015639 | 0.2829 | 0.00036959 | 1 |
| Curated_gene_sets:weber_methylated_icp_in_sperm_up | 7 | 0.94313 | 0.018276 | 0.28352 | 0.00044074 | 1 |
| Curated_gene_sets:nikolsky_breast_cancer_12q13_q21_amplicon | 45 | 0.57336 | 0.028141 | 0.1763 | 0.00057368 | 1 |
| GO_mf:go_insulin_receptor_binding | 30 | 0.44713 | 0.017926 | 0.14047 | 0.00072997 | 1 |
| GO_bp:go_positive_regulation_of_execution_phase_of_apoptosis | 11 | 0.71149 | 0.017281 | 0.22986 | 0.0009847 | 1 |
| GO_bp:go_diencephalon_development | 69 | 0.30331 | 0.018422 | 0.099388 | 0.0011391 | 1 |
| Curated_gene_sets:whitfield_cell_cycle_g2 | 171 | 0.17555 | 0.016739 | 0.05822 | 0.0012854 | 1 |

#### *5. Results of the phenotype Lack of Behavioral Control (LBCN)*

Manhattan and Q-Q plots of SNP data, Manhattan and Q-Q plots of gene-based data (MAGMA) and the results of gene-property tissue-specific expression analysis (MAGMA; GTEx v6, 53 tissue types) are shown in Figures SI5-SI9. Figure SI10 shows adjusted $R^2$s of PRS for neuroticism at different p-value thresholds on the residuals of a model regressing the personality dimension LBCN onto a set of baseline variables (see Methods and Materials). Top 10 GWAS SNPs are listed in Table SI3, Table SI4 shows the top 10 associated gene-sets from MAGMA gene-set analysis.

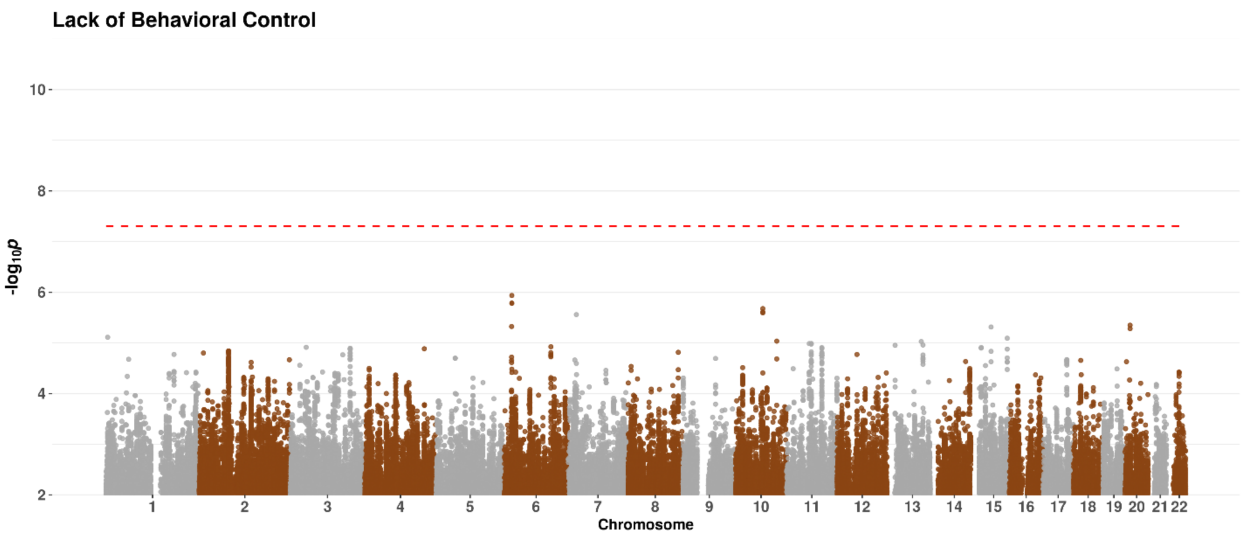


Figure SI5. Manhattan plot of the phenotype LBCN. Genome-wide significance level is indicated by the red dashed line.

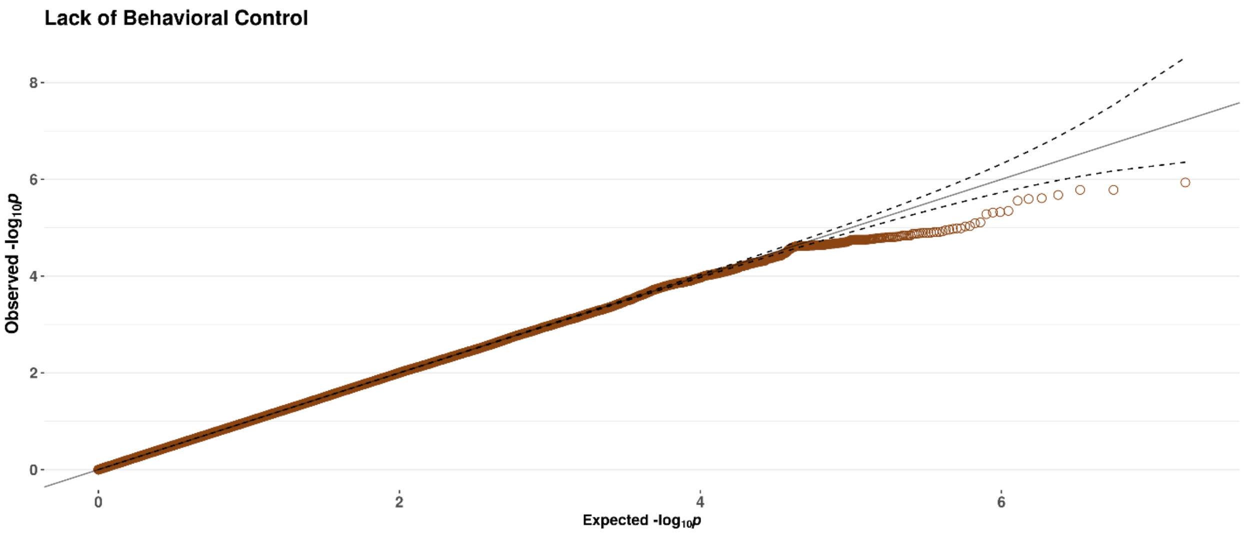


Figure SI6. Q-Q plot of the phenotype LBCN (λ=1.023).

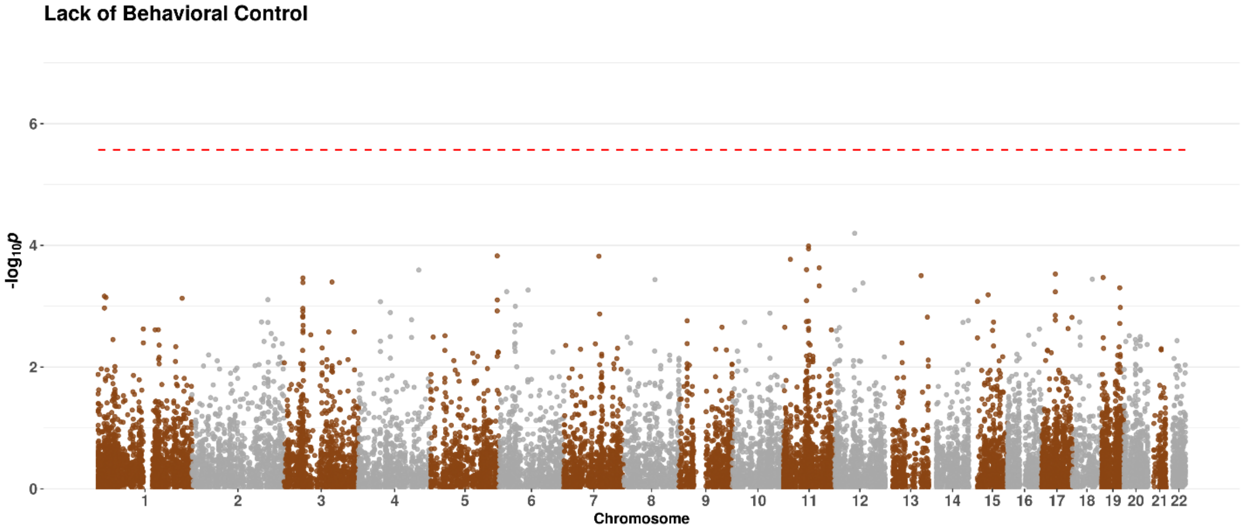


Figure SI7. Manhattan plot of the phenotype LBCN (gene-based test). Genome-wide significance level (Bonferroni-corrected for 18 634 genes) is indicated by the red dashed line.

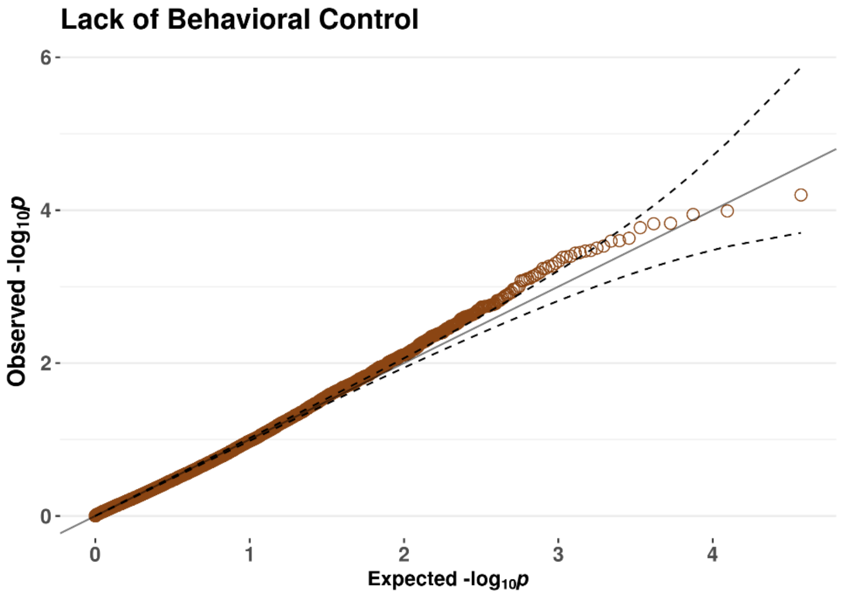


Figure SI8. Q-Q plot of the phenotype LBCN (gene-based test).

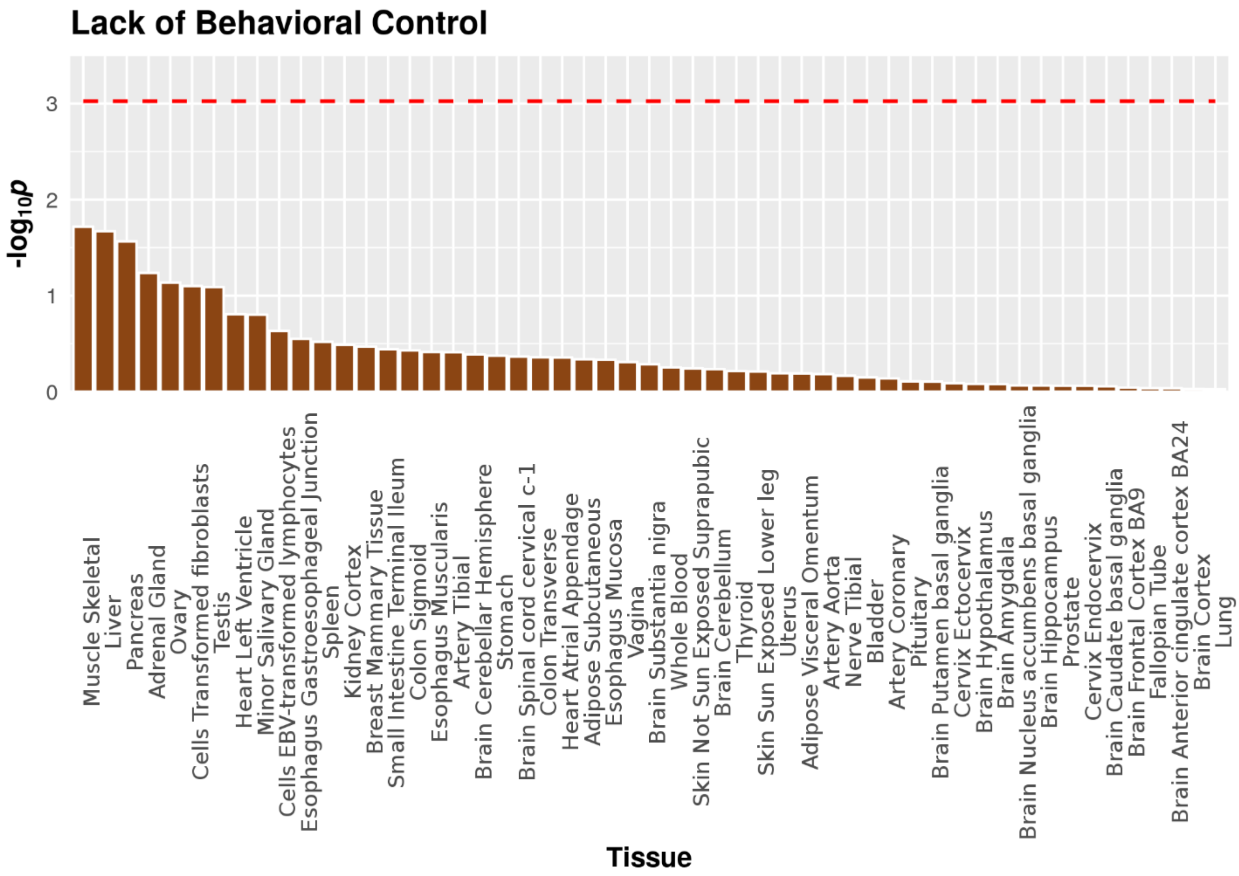


Figure SI9. Results of MAGMA gene-property tissue expression analysis of the phenotype LBCN. Bonferroni-corrected significance level (for 53 tissues) is indicated by the red dashed line.

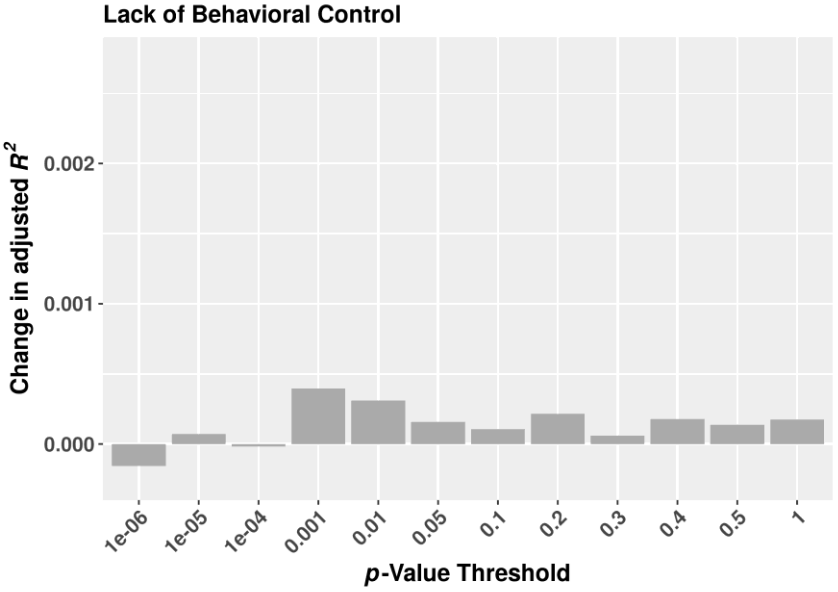


Figure SI10. Effects (adjusted $R^2$s) of PRS for neuroticism at different p-value thresholds on the residuals of a model regressing the personality dimension LBCN onto a set of baseline variables (see Methods and Materials). The range of the y-axis matches Figure 5.

Table SI3. Top ten SNPs from the GWAS of the phenotype LBCN (meta-analysis of both HeiDE samples). For abbreviations see Table SI1.

| MarkerName | Chr | Position | Allele1 | Allele2 | Freq1 | FreqSE | Effect | SE | p-value | Direction |
|---|---|---|---|---|---|---|---|---|---|---|
| rs9942459 | 6 | 20730801 | t | c | 0.6789 | 0.0014 | 0.1297 | 0.0267 | 1.162e-06 | ++ |
| rs12197544 | 6 | 20730815 | t | c | 0.6781 | 0.0013 | 0.1274 | 0.0266 | 1.652e-06 | ++ |
| rs9942429 | 6 | 20730808 | a | g | 0.322 | 0.0012 | -0.1274 | 0.0266 | 1.652e-06 | -- |
| rs185189901 | 10 | 72369449 | a | g | 0.9836 | 0 | -0.5206 | 0.1098 | 2.105e-06 | -? |
| rs117924707 | 10 | 72392744 | a | g | 0.9834 | 0 | -0.5148 | 0.1093 | 2.455e-06 | -? |
| rs184158613 | 10 | 72397071 | a | g | 0.0166 | 0 | 0.5135 | 0.1092 | 2.548e-06 | +? |
| rs144185241 | 7 | 21876685 | g | gatatta | 0.0699 | 0.0062 | -0.2321 | 0.0495 | 2.772e-06 | -- |
| rs2876363 | 20 | 13808426 | t | g | 0.1615 | 0.0042 | -0.1481 | 0.0323 | 4.484e-06 | -- |
| rs2479091 | 6 | 20142872 | c | g | 0.7846 | 0.0036 | -0.1393 | 0.0304 | 4.753e-06 | -- |
| rs56670848 | 15 | 54204953 | g | gc | 0.2819 | 0.0073 | 0.1266 | 0.0277 | 4.871e-06 | ++ |

Table SI4. Top 10 gene-sets associated with the phenotype LBCN (meta-analysis of both HeiDE samples). For abbreviations see Table SI2.

| Gene Set | NGenes | Beta | Beta STD | SE | *p*-value | $p_{Bon}$ |
|---|---|---|---|---|---|---|
| Curated_gene_sets:li_amplified_in_lung_cancer | 165 | 0.24408 | 0.022866 | 0.058148 | 1.3554e-05 | 0.144661842 |
| GO_bp:go_strand_displacement | 26 | 0.58339 | 0.021776 | 0.15072 | 5.4461e-05 | 0.581207792 |
| GO_bp:go_regulation_of_odontogenesis | 21 | 0.58453 | 0.019611 | 0.17276 | 0.00035859 | 1 |
| GO_bp:go_cell_recognition | 123 | 0.25027 | 0.020266 | 0.075163 | 0.00043564 | 1 |
| Curated_gene_sets:ikeda_mir1_targets_dn | 7 | 1.1858 | 0.022977 | 0.35673 | 0.00044462 | 1 |
| GO_bp:go_positive_regulation_of_fibroblast_migration | 10 | 0.80083 | 0.018546 | 0.24272 | 0.0004855 | 1 |
| GO_bp:go_rrna_metabolic_process | 238 | 0.15001 | 0.016845 | 0.045484 | 0.0004877 | 1 |
| Curated_gene_sets:ma_pituitary_fetal_vs_adult_dn | 19 | 0.55419 | 0.017687 | 0.16931 | 0.00053287 | 1 |
| Curated_gene_sets:matzuk_early_antral_follicle | 12 | 0.88673 | 0.022495 | 0.27743 | 0.00069749 | 1 |
| Curated_gene_sets:mikkelsen_dedifferentiated_state_dn | 7 | 0.9362 | 0.018141 | 0.30064 | 0.00092439 | 1 |

### *6. Results of the phenotype Type-A-Behavior (TYAB)*

Manhattan and Q-Q plots of SNP data, and the results of gene-property tissue-specific expression analysis (MAGMA; GTEx v6, 53 tissue types) are shown in Figures SI11-SI13. Figure SI14 shows adjusted $R^2$s of PRS for neuroticism at different p-value thresholds on the residuals of a model regressing the personality dimension TYAB onto a set of baseline variables (see Methods and Materials). Top 10 GWAS SNPs are listed in Table SI5, Table SI6 shows the top 10 associated gene-sets from MAGMA gene-set analysis.

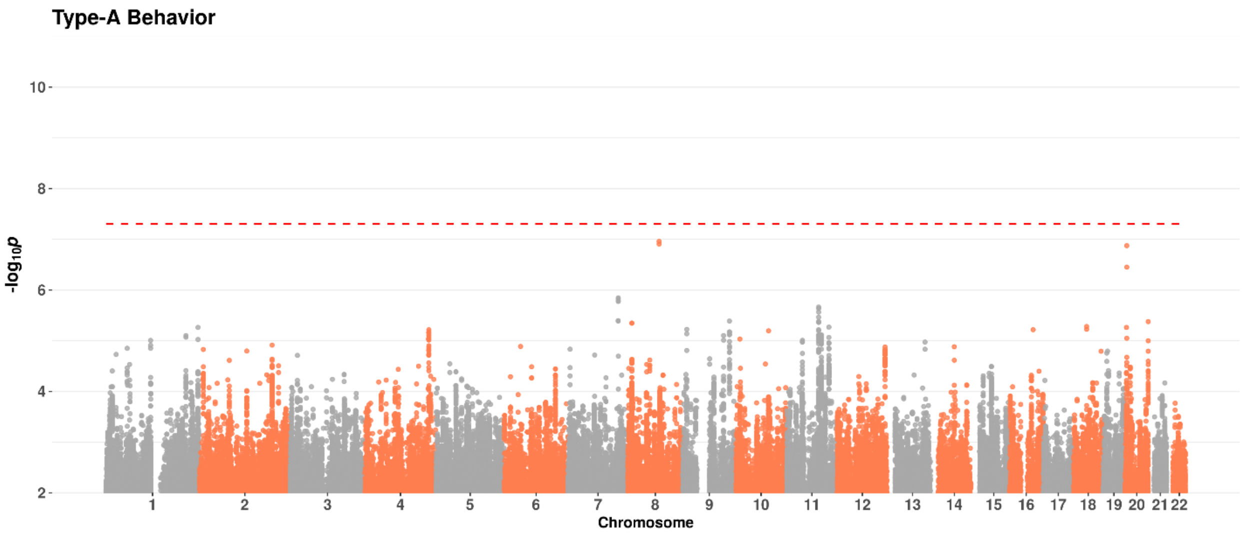


Figure SI11. Manhattan plot of the phenotype TYAB. Genome-wide significance level is indicated by the red dashed line.

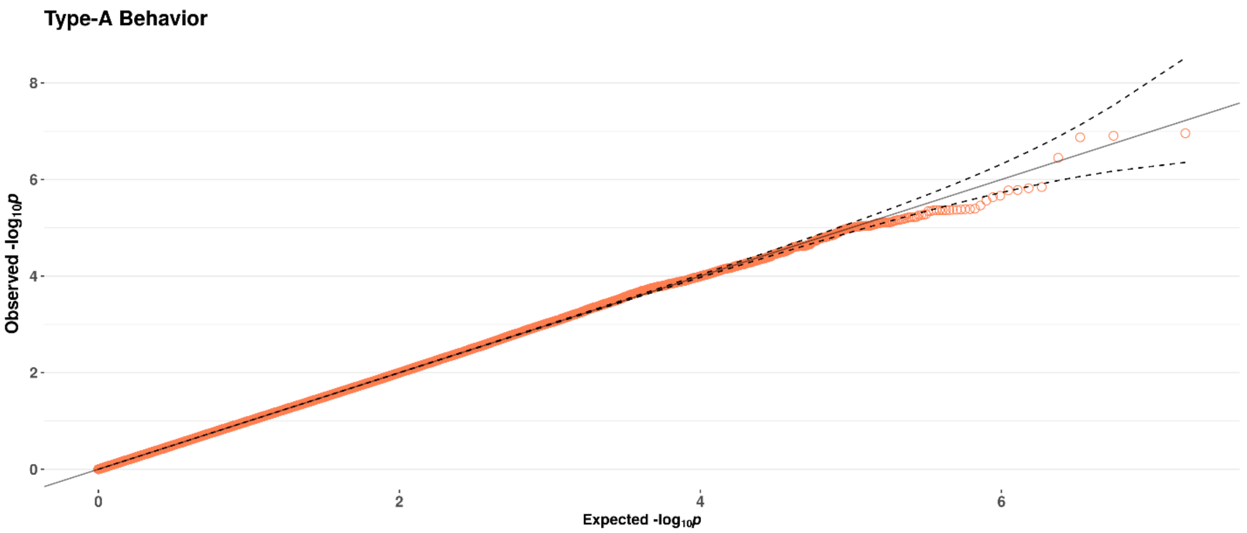


Figure SI12. Q-Q plot of the phenotype TYAB (λ=1.011).

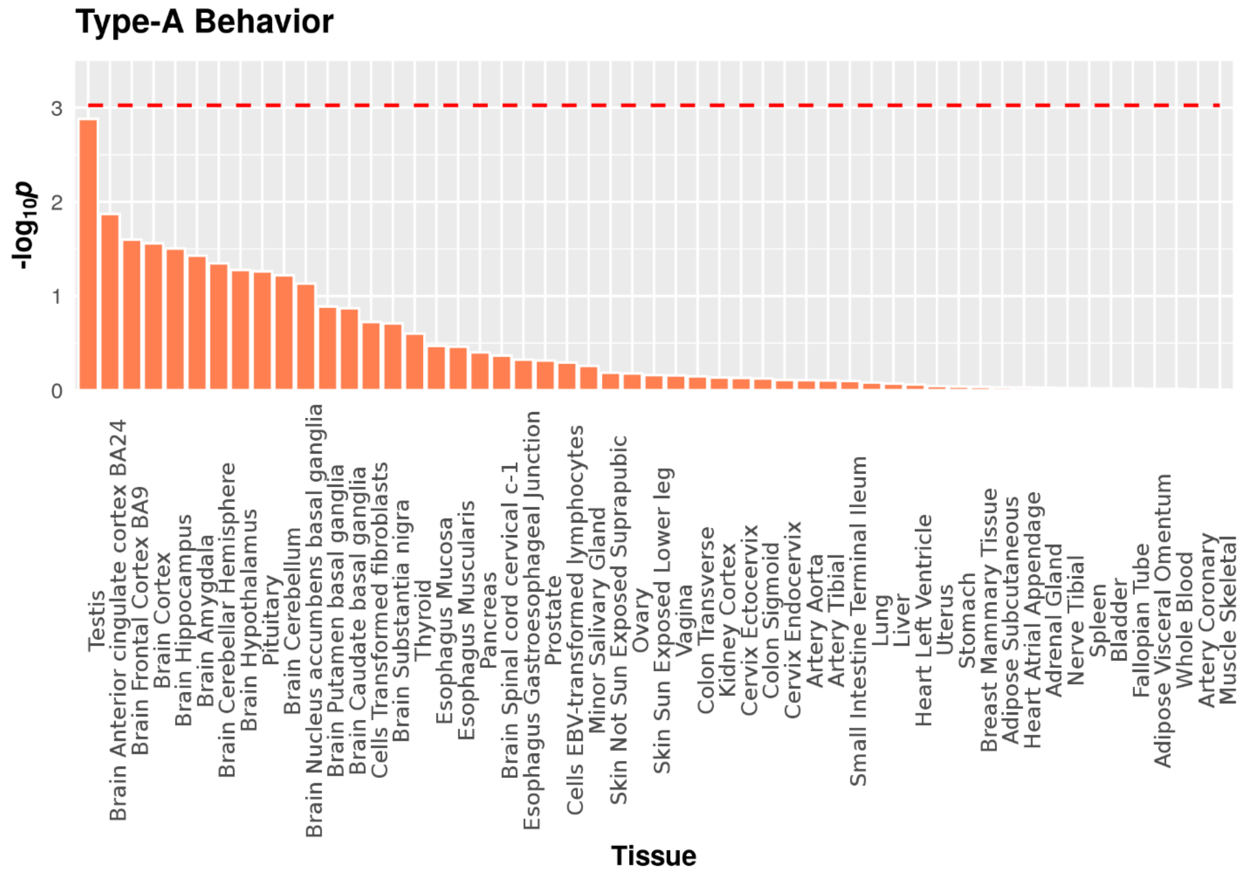


Figure SI13. Results of MAGMA gene-property tissue expression analysis of the phenotype TYAB. Bonferroni-corrected significance level (for 53 tissues) is indicated by the red dashed line.

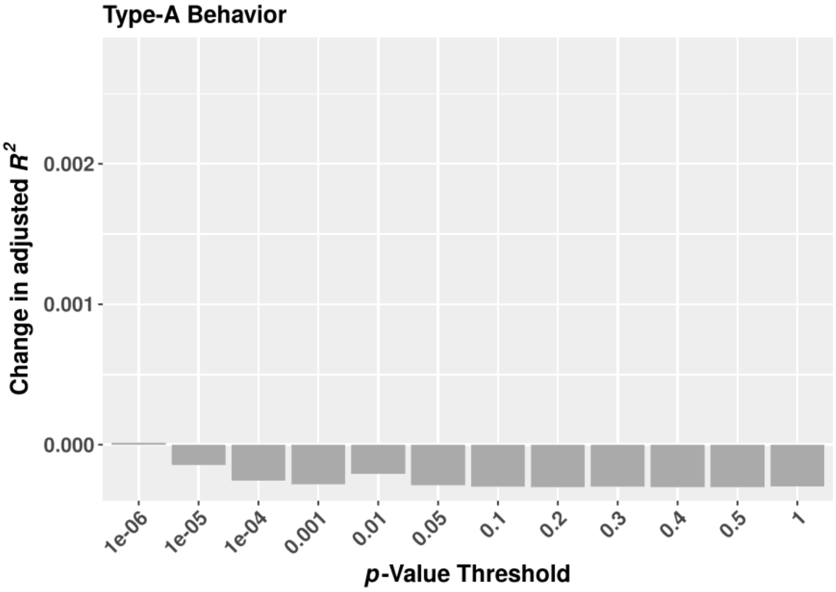


Figure SI14. Effects (adjusted $R^2$s) of PRS for neuroticism at different p-value thresholds on the residuals of a model regressing the personality dimension TYAB onto a set of baseline variables (see Methods and Materials). The range of the y-axis matches Figure 5.

Table SI5. Top ten SNPs from the GWAS of the phenotype TYAB (meta-analysis of both HeiDE samples). For abbreviations see Table SI1.

| MarkerName | Chr | Position | Allele1 | Allele2 | Freq1 | FreqSE | Effect | SE | *p*-value | Direction |
|---|---|---|---|---|---|---|---|---|---|---|
| rs142694624 | 8 | 83094903 | t | c | 0.0105 | 0 | 1.3006 | 0.245 | 1.103e-07 | ?+ |
| rs117898492 | 8 | 83250574 | t | c | 0.0105 | 0 | 1.2984 | 0.2456 | 1.244e-07 | ?+ |
| rs6084912 | 20 | 4826657 | a | c | 0.1623 | 0.0067 | -0.1906 | 0.0361 | 1.339e-07 | -- |
| rs35452355 | 20 | 4826252 | t | c | 0.7836 | 0.0126 | 0.1607 | 0.0316 | 3.552e-07 | ++ |
| rs137925278 | 7 | 133496882 | a | c | 0.9897 | 2e-04 | -0.6231 | 0.1293 | 1.433e-06 | -- |
| rs144521884 | 7 | 133515512 | c | g | 0.9898 | 2e-04 | -0.6217 | 0.1293 | 1.517e-06 | -- |
| rs146768392 | 7 | 133540427 | t | c | 0.0102 | 2e-04 | 0.6193 | 0.1292 | 1.653e-06 | ++ |
| rs142297540 | 7 | 133542283 | t | c | 0.0102 | 2e-04 | 0.6188 | 0.1292 | 1.681e-06 | ++ |
| rs56160063 | 11 | 85596214 | a | g | 0.2459 | 0.0044 | 0.1325 | 0.028 | 2.174e-06 | ++ |
| rs35944027 | 11 | 85590667 | a | at | 0.7467 | 0.0057 | -0.1331 | 0.0282 | 2.323e-06 | -- |

Table SI6. Top 10 gene-sets associated with the phenotype TYAB (meta-analysis of both HeiDE samples). For abbreviations see Table SI2.

| Gene Set | NGenes | Beta | Beta STD | SE | *p*-value | $p_{Bon}$ |
|---|---|---|---|---|---|---|
| Curated_gene_sets:inamura_lung_cancer_scc_subtypes_up | 15 | 0.81523 | 0.02312 | 0.20994 | 5.1771e-05 | 0.552551883 |
| GO_bp:go_negative_regulation_of_jun_kinase_activity | 14 | 0.84861 | 0.023251 | 0.22522 | 8.2609e-05 | 0.881603248 |
| GO_bp:go_negative_regulation_of_jnk_cascade | 33 | 0.48267 | 0.020294 | 0.13243 | 0.00013431 | 1 |
| GO_bp:go_negative_reg_of_stress_act_prot_kin_sign_cas | 40 | 0.43914 | 0.020323 | 0.12332 | 0.00018528 | 1 |
| GO_mf:go_myosin_binding | 56 | 0.33875 | 0.018542 | 0.099589 | 0.00033595 | 1 |
| Curated_gene_sets:wang_lmo4_targets_dn | 330 | 0.14446 | 0.019053 | 0.043001 | 0.00039145 | 1 |
| Curated_gene_sets:weber_methylated_hcp_in_fibroblast_dn | 38 | 0.41619 | 0.018775 | 0.13055 | 0.0007175 | 1 |
| GO_mf:go_heparan_sulfate_proteoglycan_binding | 16 | 0.62952 | 0.018438 | 0.19768 | 0.00072608 | 1 |
| Curated_gene_sets:krasnoselskaya_ilf3_targets_dn | 43 | 0.3824 | 0.018348 | 0.12155 | 0.00082936 | 1 |
| GO_cc:go_prp19_complex | 13 | 0.58405 | 0.015421 | 0.18637 | 0.00086432 | 1 |

### *7. Results of the phenotype Locus of Control over Disease (LOCC)*

Manhattan and Q-Q plots of SNP data, Manhattan and Q-Q plots of gene-based data (MAGMA) and the results of gene-property tissue-specific expression analysis (MAGMA; GTEx v6, 53 tissue types) are shown in Figures SI15-SI19. Figure SI20 shows adjusted $R^2$s of PRS for neuroticism at different p-value thresholds on the residuals of a model regressing the personality dimension LOCC onto a set of baseline variables (see Methods and Materials). Top 10 GWAS SNPs are listed in Table SI7, Table SI8 shows the top 10 associated gene-sets from MAGMA gene-set analysis.

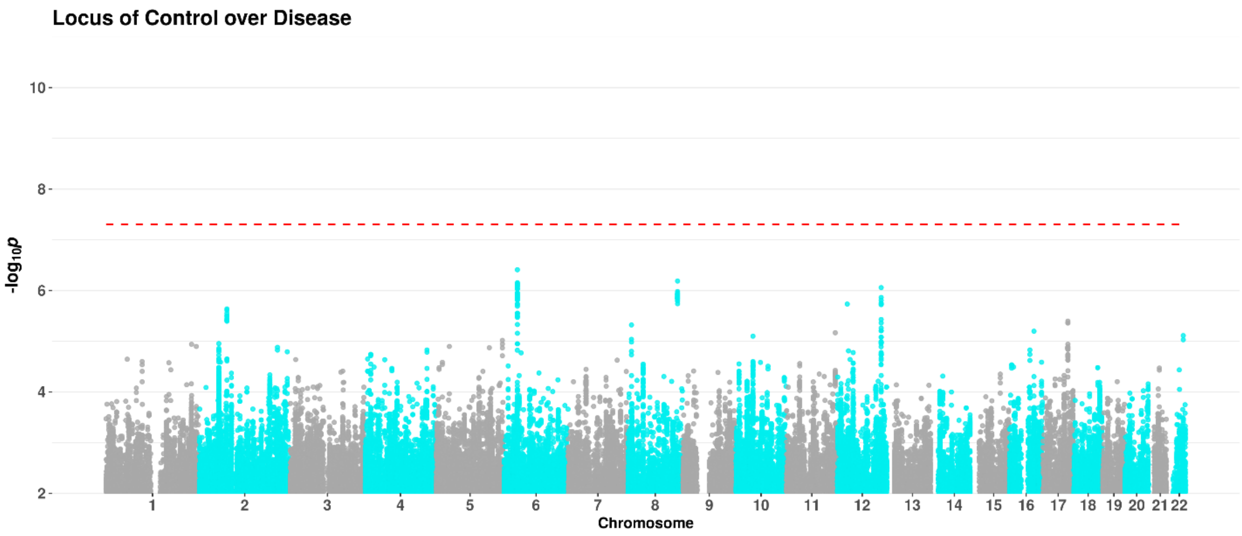


Figure SI15. Manhattan plot of the phenotype LOCC. Genome-wide significance level is indicated by the red dashed line.

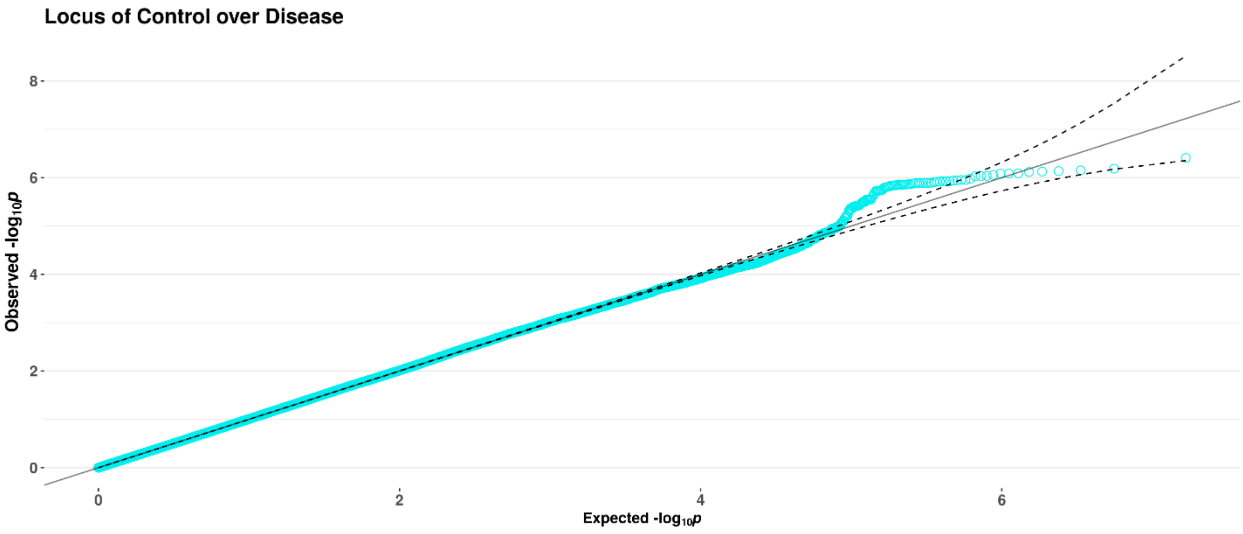


Figure SI16. Q-Q plot of the phenotype LOCC (λ= 1.004).

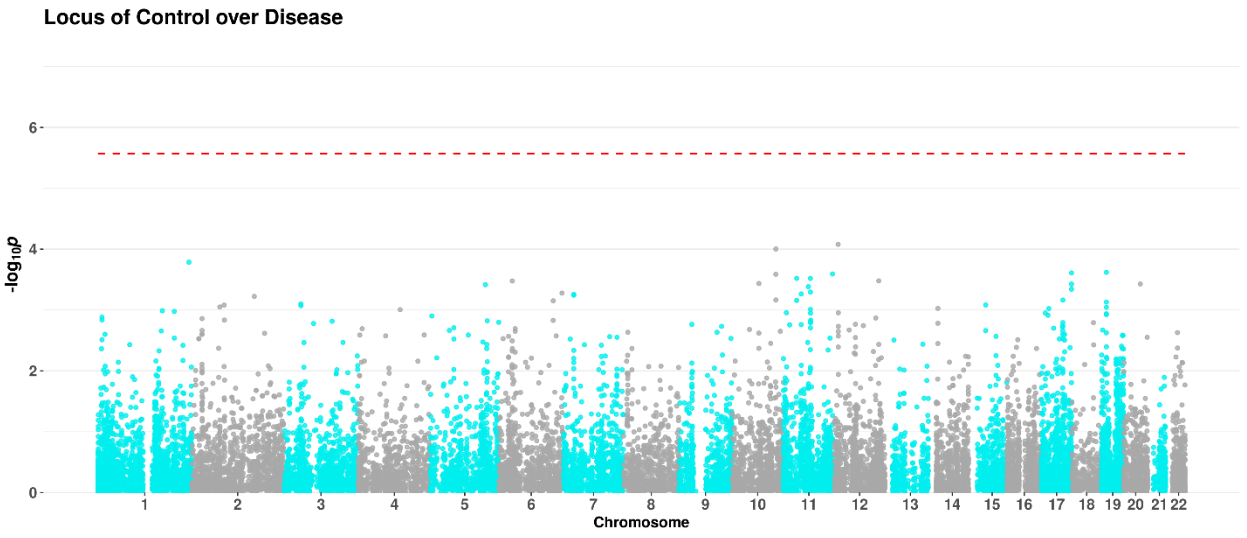


Figure SI17. Manhattan plot of the phenotype LOCC (gene-based test). Genome-wide significance level (Bonferroni-corrected for 18 634 genes) is indicated by the red dashed line.

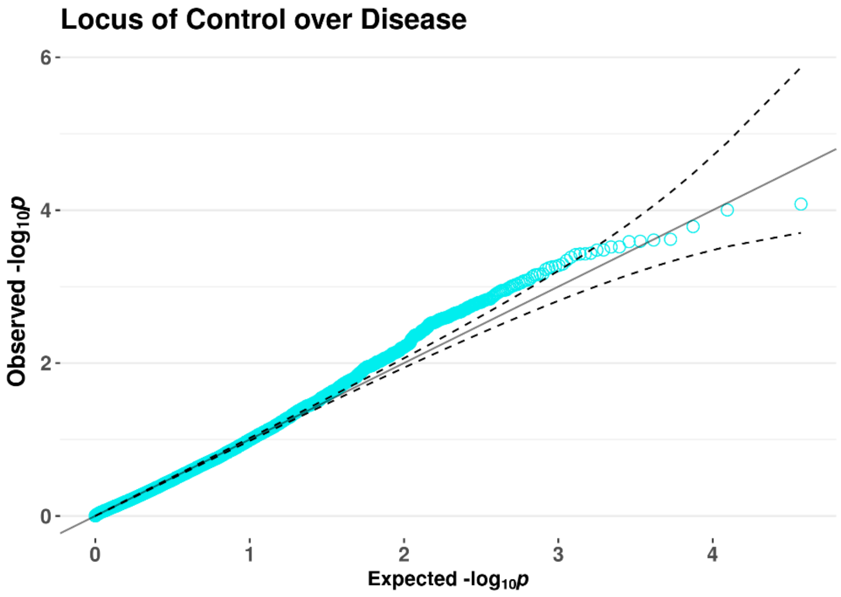


Figure SI18. Q-Q plot of the phenotype LOCC (gene-based test).

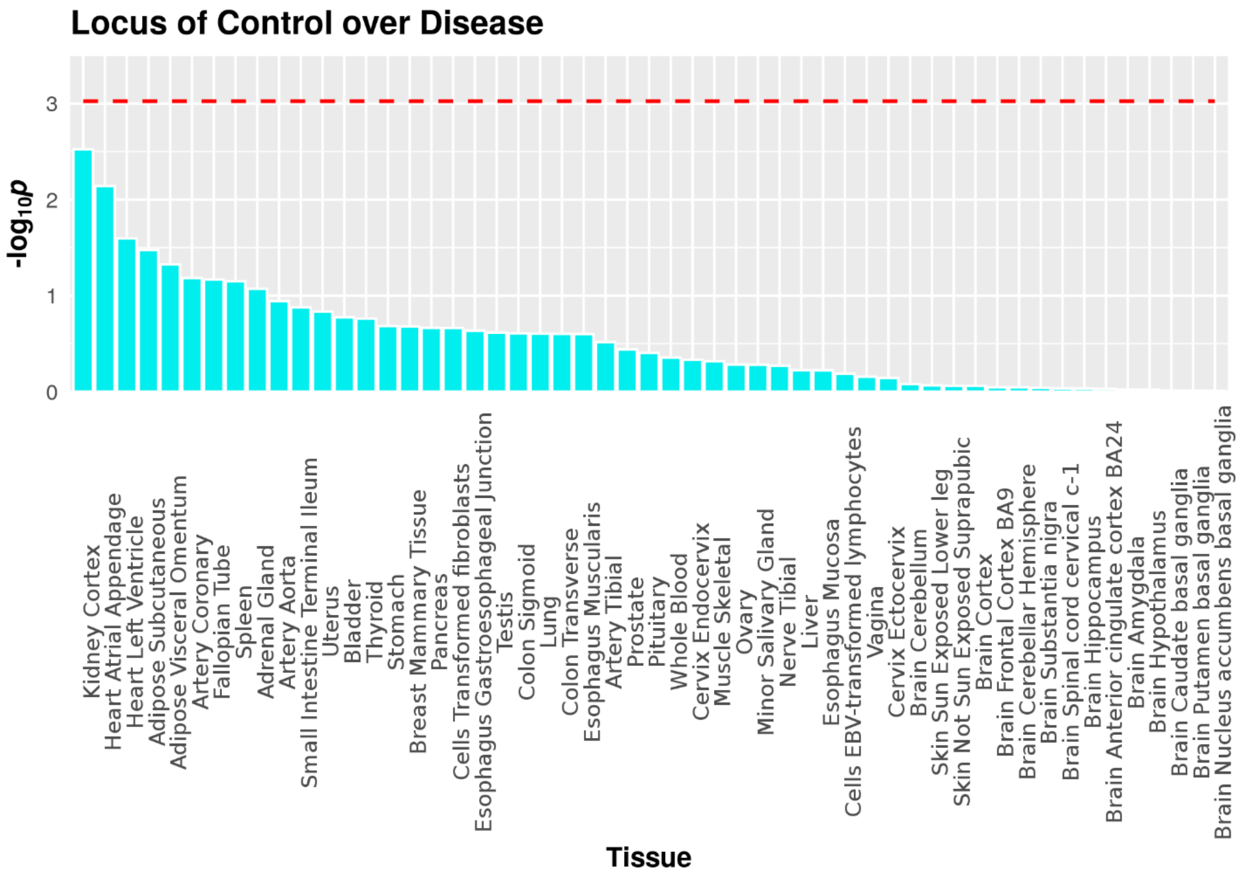


Figure SI19. Results of MAGMA gene-property tissue expression analysis of the phenotype LOCC. Bonferroni-corrected significance level (for 53 tissues) is indicated by the red dashed line.

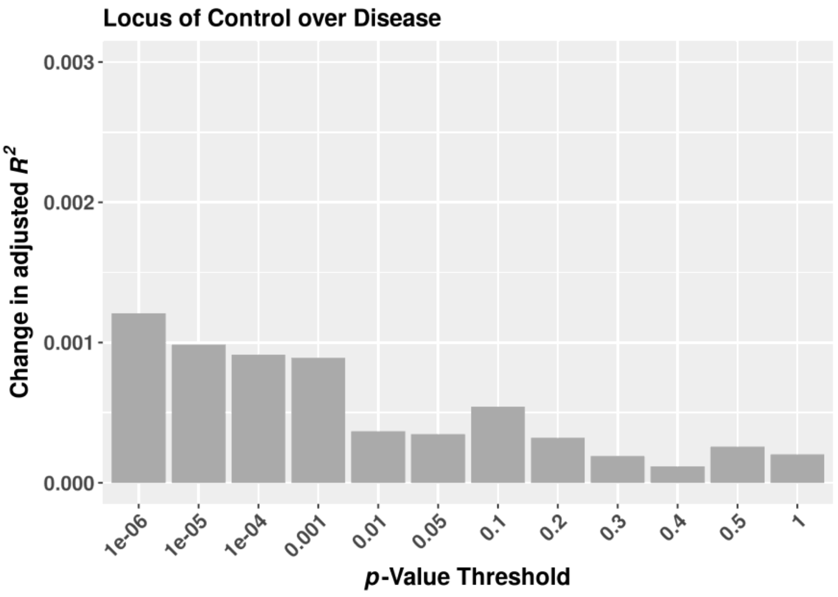


Figure SI20. Effects (adjusted $R^2$s) of PRS for neuroticism at different p-value thresholds on the residuals of a model regressing the personality dimension LOCC onto a set of baseline variables (see Methods and Materials). The range of the y-axis matches Figure 5.

Table SI7. Top ten SNPs from the GWAS of the phenotype LOCC (meta-analysis of both HeiDE samples). For abbreviations see Table SI1.

| MarkerName | Chr | Position | Allele1 | Allele2 | Freq1 | FreqSE | Effect | SE | *p*-value | Direction |
|---|---|---|---|---|---|---|---|---|---|---|
| 6:35732878:CA | 6 | 35732878 | ca | caa | 0.3843 | 0.0047 | -0.1262 | 0.0249 | 3.906e-07 | -- |
| rs1812948 | 8 | 132581723 | a | g | 0.1975 | 0.0076 | -0.1525 | 0.0306 | 6.508e-07 | -- |
| rs9470089 | 6 | 35733305 | a | g | 0.3811 | 0.0045 | -0.1231 | 0.0248 | 7.043e-07 | -- |
| rs2817054 | 6 | 35733699 | a | g | 0.3779 | 0.0049 | -0.1229 | 0.0248 | 7.31e-07 | -- |
| rs2817053 | 6 | 35733461 | a | g | 0.3779 | 0.0049 | -0.1229 | 0.0248 | 7.466e-07 | -- |
| rs2766567 | 6 | 35734094 | a | g | 0.377 | 0.0049 | -0.1231 | 0.0249 | 7.618e-07 | -- |
| rs2817055 | 6 | 35733778 | c | g | 0.6227 | 0.0047 | 0.1226 | 0.0248 | 8.083e-07 | ++ |
| rs35662542 | 6 | 35734704 | ca | c | 0.6194 | 0.0041 | 0.1225 | 0.0248 | 8.171e-07 | ++ |
| rs2817057 | 6 | 35734884 | a | g | 0.3807 | 0.0041 | -0.1225 | 0.0248 | 8.262e-07 | -- |
| rs3782193 | 12 | 117497066 | a | c | 0.1508 | 0.0035 | 0.1664 | 0.0338 | 8.792e-07 | ++ |

Table SI8. Top 10 gene-sets associated with the phenotype LOCC (meta-analysis of both HeiDE samples). For abbreviations see Table SI2.

| Gene Set | NGenes | Beta | Beta STD | SE | *p*-value | $p_{Bon}$ |
|---|---|---|---|---|---|---|
| GO_bp:go_cellular_response_to_ionizing_radiation | 51 | 0.50168 | 0.026209 | 0.10925 | 2.2102e-06 | 0.0235894646 |
| GO_bp:go_cardiac_ventricle_development | 104 | 0.327 | 0.02436 | 0.082079 | 3.4037e-05 | 0.363242864 |
| GO_bp:go_circulatory_system_development | 756 | 0.11653 | 0.02299 | 0.029925 | 4.948e-05 | 0.52800108 |
| GO_bp:go_response_to_ionizing_radiation | 141 | 0.25784 | 0.022343 | 0.066488 | 5.2865e-05 | 0.56406955 |
| GO_bp:go_cardiac_muscle_tissue_development | 137 | 0.25825 | 0.022061 | 0.066955 | 5.7599e-05 | 0.614523731 |
| Curated_gene_sets:kim_wt1_targets_up | 212 | 0.21678 | 0.02299 | 0.057622 | 8.4528e-05 | 0.901744704 |
| GO_bp:go_cellular_response_to_radiation | 132 | 0.24994 | 0.020961 | 0.067445 | 0.0001057 | 1 |
| Curated_gene_sets:reactome_signaling_by_fgfr3_m | 11 | 0.94905 | 0.023051 | 0.26207 | 0.00014699 | 1 |
| GO_bp:go_morphogenesis_of_embryonic_epithelium | 131 | 0.25016 | 0.020901 | 0.071939 | 0.00025378 | 1 |
| GO_bp:go_cardiac_ventricle_morphogenesis | 60 | 0.37158 | 0.02105 | 0.10726 | 0.00026666 | 1 |

#### *8. Results of the phenotype Psychoticism (PSYC)*

Figure SI21 shows adjusted $R^2$s of PRS for neuroticism at different p-value thresholds on the residuals of a model regressing the personality dimension PSYC onto a set of baseline variables (see Methods and Materials). Table SI9 shows the top 10 associated gene-sets from MAGMA gene-set analysis.

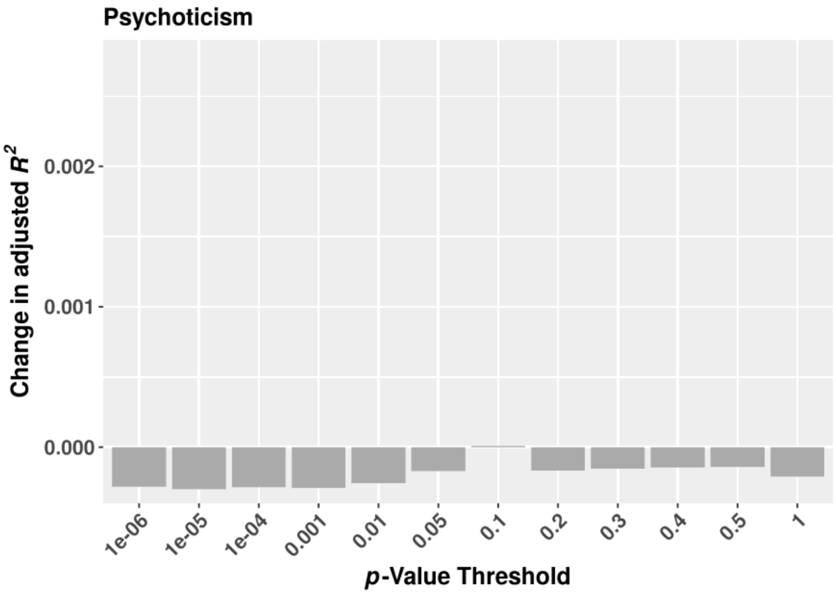


Figure SI21. Effects (adjusted $R^2$s) of PRS for neuroticism at different p-value thresholds on the residuals of a model regressing the personality dimension PSYC onto a set of baseline variables (see Methods and Materials). The range of the y-axis matches Figure 5.

Table SI9. Top 10 gene-sets associated with the phenotype PSYC (meta-analysis of both HeiDE samples). For abbreviations see Table SI2.

| Gene Set | NGenes | Beta | Beta STD | SE | *p*-value | $p_{Bon}$ |
|---|---|---|---|---|---|---|
| Curated_gene_sets:biocarta_g1_pathway | 28 | 0.63367 | 0.024544 | 0.15049 | 1.2802e-05 | 0.136635746 |
| GO_bp:go_cell_projection_organization | 853 | 0.10905 | 0.02279 | 0.028151 | 5.3816e-05 | 0.574324352 |
| GO_mf:go_c_c_chemokine_receptor_activity | 12 | 0.84699 | 0.021486 | 0.23229 | 0.00013348 | 1 |
| Curated_gene_sets:liu_common_cancer_genes | 73 | 0.31973 | 0.019972 | 0.090217 | 0.0001976 | 1 |
| Curated_gene_sets:kegg_prostate_cancer | 84 | 0.30342 | 0.020325 | 0.087132 | 0.00024921 | 1 |
| Curated_gene_sets:reactome_g1_s_spec_transcr | 16 | 0.58815 | 0.017227 | 0.17343 | 0.00034859 | 1 |
| Curated_gene_sets:browne_hcmv_inf_20hr_dn | 97 | 0.2765 | 0.019897 | 0.082619 | 0.00040979 | 1 |
| GO_bp:go_epithelial_cell_apoptotic_process | 25 | 0.54042 | 0.019781 | 0.16159 | 0.00041326 | 1 |
| GO_bp:go_syncytium_formation | 25 | 0.55458 | 0.020299 | 0.17135 | 0.00060629 | 1 |
| Curated_gene_sets:lee_liver_cancer_myc_dn | 59 | 0.36665 | 0.020598 | 0.11384 | 0.00064061 | 1 |

***9. Follow-up analyses: English translation of screening questions for anxiety disorders at the second HeiDE follow-up (2013)***

1. “Have you ever suffered for several months from bodily discomfort, for which your doctor had no conclusive explanation?”

2. “Have you ever experienced an anxiety or panic attack, during which you were overwhelmed by a feeling of strong anxiety, unease or unrest?”

3. “In your life, was there ever a time of one month (or longer) in which you were felt anxious, tense or worried often or most of the time?”

4. “Have you ever suffered from unfounded anxiety in social situations such as talking to others, doing something in front of others or being the center of attention?”

5. “Did you ever suffer from unfounded anxiety of using public transportation, going into shops or staying in public places”?

6. “Was there ever a time in which you suffered from unfounded anxiety of other situations (e.g. closed spaces) or things (e.g. heights, thunderstorms or animals)?”

***10. Diagnostic plots of the longitudinal regression analyses of depressive and anxiety symptoms***

Using the R autoplot package, we examined several diagnostic plots, including Residuals vs. Leverage plots (see below), and decided to include all individuals in the regression analyses.

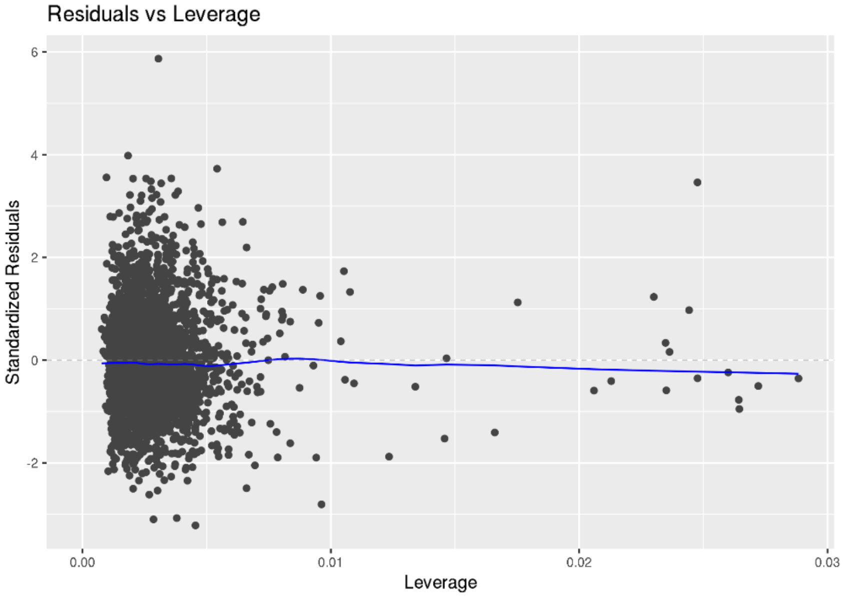


Figure SI 22. Residuals vs. Leverage plot of depressive symptoms.

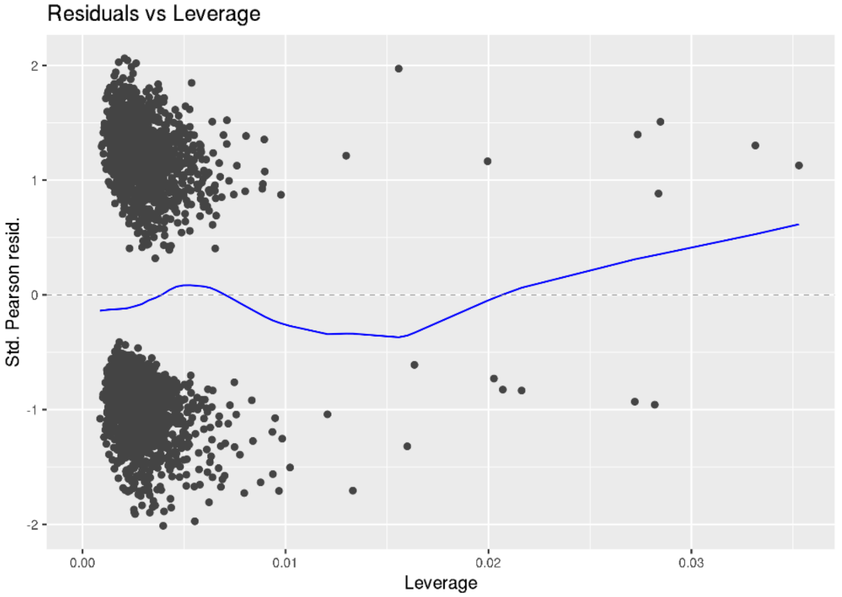


Figure SI 23. Residuals vs. Leverage plot of anxiety symptoms.
